# A public antibody class recognizes a novel S2 epitope exposed on open conformations of SARS-CoV-2 spike

**DOI:** 10.1101/2021.12.01.470767

**Authors:** Mathieu Claireaux, Tom G Caniels, Marlon de Gast, Julianna Han, Denise Guerra, Gius Kerster, Barbera DC van Schaik, Aldo Jongejan, Angela I. Schriek, Marloes Grobben, Philip JM Brouwer, Karlijn van der Straten, Yoann Aldon, Joan Capella-Pujol, Jonne L Snitselaar, Wouter Olijhoek, Aafke Aartse, Mitch Brinkkemper, Ilja Bontjer, Judith A Burger, Meliawati Poniman, Tom PL Bijl, Jonathan L Torres, Jeffrey Copps, Isabel Cuella Martin, Steven W de Taeye, Godelieve J de Bree, Andrew B Ward, Kwinten Sliepen, Antoine HC van Kampen, Perry D Moerland, Rogier W Sanders, Marit J van Gils

**Author notes:** Corresponding authors. (R.W.S.); (M.J.v.G.). These authors contributed equally (Mathieu Claireaux, Tom G Caniels).

## Abstract

Delineating the origins and properties of antibodies elicited by SARS-CoV-2 infection and vaccination is critical for understanding their benefits and potential shortcomings. Therefore, we investigated the SARS-CoV-2 spike (S)-reactive B cell repertoire in unexposed individuals by flow cytometry and single-cell sequencing. We found that ∼82% of SARS-CoV-2 S-reactive B cells show a naive phenotype, which represents an unusually high fraction of total human naive B cells (∼0.1%). Approximately 10% of these naive S-reactive B cells shared an IGHV1-69/IGKV3-11 B cell receptor pairing, an enrichment of 18-fold compared to the complete naive repertoire. A proportion of memory B cells, comprising switched (∼0.05%) and unswitched B cells (∼0.04%), was also reactive with S and some of these cells were reactive with ADAMTS13, which is associated with thrombotic thrombocytopenia. Following SARS-CoV-2 infection, we report an average 37-fold enrichment of IGHV1-69/IGKV3-11 B cell receptor pairing in the S-reactive memory B cells compared to the unselected memory repertoire. This class of B cells targets a previously undefined non-neutralizing epitope on the S2 subunit that becomes exposed on S proteins used in approved vaccines when they transition away from the native pre-fusion state because of instability. These findings can help guide the improvement of SARS-CoV-2 vaccines.

## Introduction

The emergence of severe acute respiratory syndrome coronavirus 2 (SARS-CoV-2) at the end of 2019 and its increased spread as a result of novel viral variants has posed considerable danger to global health. Multiple vaccines are now in use that confer high levels of protection. As of November 25th 2021, 54% of the world’s population has received at least one dose of a coronavirus disease 2019 (COVID-19) vaccine, illustrating the rapid development and distribution of vaccines (Ritchie et al. 2020). A major goal of these vaccines is to induce neutralizing antibodies (NAbs) and memory B cells that protect against subsequent infection.

Most licensed vaccines aim to induce immunity against SARS-CoV-2 S, a trimeric glycoprotein on the surface of the virion, that is the only known target for NAbs. It consists of an apical S1 subunit encompassing an N-terminal domain (NTD) and the receptor-binding domain (RBD) that is responsible for binding to the ACE2 receptor; and a membrane-proximal S2 subunit which is responsible for fusion of viral and cellular membranes. Coronavirus (CoV) S proteins can suffer from instability and deteriorate into non-native forms leading to altered exposure of antibody epitopes (Hsieh et al. 2020; Berger and Schaffitzel 2020; Juraszek et al. 2021). Therefore, some of the approved vaccines, including those from J&J/Janssen, Moderna and Pfizer/BioNTech, but not those from Oxford/AstraZeneca and several others, contain modifications to stabilize the S proteins. These changes include the removal of the furin cleavage site between S1 and S2 and two proline substitutions (K986P/V987P) in S2, resulting in a prefusion stabilized S trimer termed S-2P (Sadoff et al. 2021; Baden et al. 2021; Polack et al. 2020). Although S-2P is more stable than wild-type S (S-WT), subsequent studies revealed that even S-2P suffers from instability issues and displays open conformations (Juraszek et al. 2021; Henderson et al. 2020; Hsieh et al. 2020). Several additional stabilization strategies have been described, including the introduction of an additional four prolines (F817P/A892P/A899P/A942P), resulting in S-6P (HexaPro S), which shows considerably increased stability and resistance to heat/freeze cycles compared to S-2P and S-WT (Hsieh et al. 2020; Henderson et al. 2020; McCallum et al. 2020; Juraszek et al. 2021).

Early in the pandemic many groups identified and isolated potent NAbs from COVID-19 patients that defined important epitopes on S, which have led to emergency use authorization of several monoclonal antibody (MAb) therapies for COVID-19 (Taylor et al. 2021). Most of the potently neutralizing MAbs target the immunodominant RBD and NTD on the apex of S, whereas MAbs against the S2 domain, while generally less potent, tend to be more broadly reactive across sarbecoviruses and endemic human CoVs (HCoVs) (Shiakolas et al. 2021; Amanat, Thapa, et al. 2021). B cells that arise during SARS-CoV-2 infection target both S1 and S2 domains, but their ontogeny is often unclear. S1 NAbs are usually poorly cross-reactive with different SARS-CoV-2 variants or other HCoVs and have not undergone extensive somatic hypermutation (SHM), suggesting that they originate from *de novo* activation of naive B cells (Kreer et al. 2020; Brouwer et al. 2020; Zost et al. 2020; Rogers et al. 2020; Robbiani et al. 2020). In contrast, a subset of S2 NAbs cross-bind and cross-neutralize endemic HCoVs, suggesting that they arise from pre-existing memory B cells, formed during infection with an endemic HCoV and reactivated upon SARS-CoV-2 infection (Song et al. 2021). Thus, it is conceivable that in SARS-CoV-2 infection and similarly, vaccination, the humoral immune response is driven by both *de novo* activation of naive B cells and reactivation of memory B cells.

Although the NAb response to SARS-CoV-2 infection and vaccination has been studied extensively, non-neutralizing MAbs (non-NAbs) have not been a focal point. While understudied, non-NAbs make up a substantial portion of the antibody repertoire after infection and the majority of vaccine-induced anti-S Abs are also non-neutralizing (Amanat, Thapa, et al. 2021; Dugan et al. 2021; Sakharkar et al. 2021). Non-NAbs can contribute to immunity through effector mechanisms such as antibody-dependent cellular cytotoxicity (ADCC) and phagocytosis (ADCP) (Ullah et al. 2021; Chan et al. 2021; Winkler et al. 2021; Schäfer et al. 2021; Shiakolas et al. 2021), while high levels of pro-inflammatory antibodies may contribute to severe disease (Chakraborty et al. 2021; Larsen et al. 2021). Non-NAbs elicited by infection target S as well as other viral proteins such as nucleoprotein (N) and ORF8 (Dugan et al. 2021), whereas the non-NAbs induced by currently approved vaccines exclusively target S. S instability can contribute to the induction of non-NAbs because their epitopes become exposed on open/aberrant conformations. Extensive work on respiratory syncytial virus (RSV) and human immunodeficiency virus 1 (HIV-1) has shown that the occurrence of post-fusion or open conformations of the surface glycoproteins leads to the induction of non-NAbs, while stabilizing these glycoproteins in the closed pre-fusion conformation reduced the induction of non-NAbs (McLellan et al. 2013; de Taeye et al. 2015; Sanders and Moore 2021). In particular for RSV, the reduced induction of non-NAbs was accompanied by an enhanced NAb response. Elucidating the properties and epitopes of non-NAbs as well as their B cell origins, is therefore highly relevant for both understanding humoral immunity against SARS-CoV-2 and improving vaccines. Here, we characterized the human baseline B cell repertoire that recognizes SARS-CoV-2 S prior to any antigenic SARS-CoV-2 S encounter through infection or vaccination. We found that an unusually high proportion (∼0.1%) of naive B cells is able to recognize SARS-CoV2 S and that these naive B cells show a highly enriched usage of heavy chain immunoglobulin V gene (IGHV)1-69 and kappa chain immunoglobulin V gene (IGKV)3-11 in their B cell receptor (BCR). A subset of these naive B cells exhibits polyreactivity and can be activated upon SARS-CoV-2 S encountering. Following SARS-CoV-2 infection this particular pairing was selected and highly enriched in the memory B cell population and almost exclusively targets a novel non-neutralizing apical S2 epitope present on aberrant forms of S, suggesting that further stabilization of S might benefit SARS-CoV-2 vaccine responses.

## Results

### Multiple B cell compartments in SARS-CoV-2 naive individuals recognize SARS-CoV-2 S

Recent studies on sera and memory B cells from recovered COVID-19 patients suggest that a fraction of the antibody response against SARS-CoV-2 S could stem from pre-existing cross-reactive memory B cells induced by prior infection with endemic HCoVs (Ng et al. 2020; Song et al. 2021; Grobben et al. 2021; Tong et al. 2021). Therefore, we interrogated the baseline SARS-CoV-2 S-reactive B cell repertoire from ten unexposed and unvaccinated healthy donors (HD01-10), sampled in 2019 and early 2020, before the first official case report of SARS-CoV-2 infection in the Netherlands. We developed a novel combinatorial B cell staining approach with labelled antigenic probes, allowing for the simultaneous identification of B cells that are reactive to six different pathogens in a single sample (Table S1). SARS-CoV-2 S-reactive B cells were compared to those recognizing pre-encountered antigens, including influenza A virus hemagglutinin (H1N1_pdm09_ HA), RSV fusion protein (RSV F) and tetanus toxoid (TT), as well as those against unencountered antigens HIV-1 envelope glycoprotein (Env) and hepatitis C virus (HCV) envelope glycoprotein (E1E2; Fig. 1A, Fig. S1A-E). B cells specific for SARS-CoV-2 were present at high frequency in unexposed individuals (median 0.086% of total B cells), which was ∼10-30-fold higher compared to the frequency of B cells reactive with other unencountered antigens HIV-1 Env and HCV E1E2 (median 0.007% and 0.0025%, respectively), but in a similar range to the frequency of B cells reactive with previously encountered antigens H1N1 HA (0.066%), RSV F (0.042%) and TT (0.12%, Fig. 1B).

**Figure 1.**
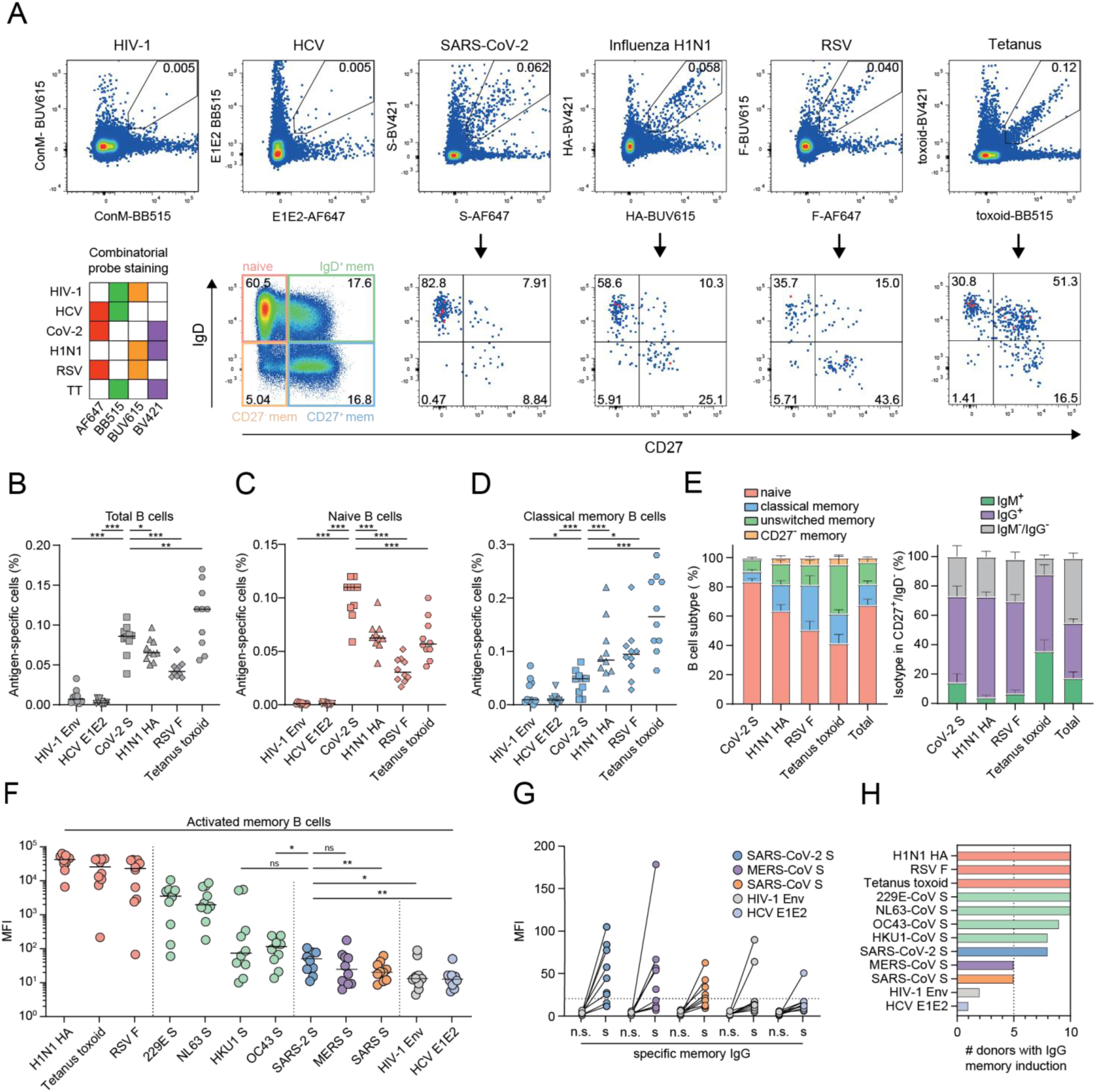
Phenotypic characterization of SARS-CoV-2 S-reactive B cells in unexposed individuals. **(A)** Combinatorial probe staining and gating strategy for the detection of multiple B cell specificities in a single PBMC sample. Top panel: From live B cells (gating strategy Fig. S1), antigen-specific B cells are detected as double positive for the binding of the same antigen multimerized with two different fluorochromes. Bottom panel left, matrix depicting each combination of two fluorochrome-coded to the same antigen. Bottom panel left, B cell subsets were determined from antigen-specific B cells according IgD and CD27 expression (IgD^+^/CD27^-^, naive; IgD^+^/CD27^+^, unswitched IgD^+^ memory; IgD^-^/CD27^-^, CD27^-^ memory; IgD^-^/CD27^+^, classical memory). The numbers inside the boxes represent the frequency (%) of cells in a gate. (**B-D**) Frequency of antigen-specific B cells for ten HDs (Statistical differences were tested only in comparison to SARS-CoV-2 condition) in total B cells (**B**), naive B cells (IgD^+^/CD27^-^, (**C**)), or memory B cells (IgD^-^/CD27^+^, (**D**)). Each dot represents one individual HD. The line represents the median frequency. (**E**) The phenotype of antigen-specific B cells for each antigen (left panel) and the isotype as detected by flow cytometry in the classical memory B cell population (IgD^-^/CD27^+^, right panel). Bars represent mean ± SD. (**F**) Median fluorescence intensity (MFI) of IgG detected in undiluted supernatants of cultured PBMCs for each antigen as measured by custom Luminex assay. Each dot represents one individual HD. (**G**) MFI of IgG detected in supernatants of cultured PBMCs as measured by Luminex assay for five antigens. n.s., not-stimulated; s., stimulated with resiquimod (TLR7/8 agonist), IL-2 and IL-21 to induce IgG secretion. The dotted line represents the cutoff of detectable IgG (three times above background). (**H**) Number of HDs that show IgG secretion specific against all antigens measured when stimulated with cytokines to induce IgG secretion. n.s., not significant; *, *p* < 0.05; **, *p* < 0.01; ***, *p* < 0.001; ****, *p* < 0.0001).

Next, we analyzed the naive and memory subsets of these antigen-specific B cells, which can be subdivided into four populations based on their surface expression of IgD and CD27: naive (IgD^+^/CD27^-^), classical memory (IgD^-^/CD27^+^), CD27^-^ memory (IgD^-^/CD27^-^) and unswitched memory (IgD^+^/CD27^+^) (Fig. 1A lower panel, C-D and Fig. S2A, B). There were virtually no naive B cells specific to HIV-1 Env and HCV E1E2 (∼0.001%) (Fig. 1C). In contrast, the proportion of B cells that recognize SARS-CoV-2 S was significantly higher (0.11%) in the naive compartment than those specific against any other probe tested (Fig. 1C). This indicates that a *de novo* B cell response stemming from the naive compartment likely plays a role in SARS-CoV-2 infection.

In the classical memory compartment, B cells specific against recurring seasonal infections (H1N1 HA and RSV F; 0.084% and 0.095%, respectively) or those elicited against a vaccine component (TT; 0.17%) were significantly more predominant than those against SARS-CoV-2 S (0.049%) (Fig. 1D), as expected. However, the SARS-CoV-2 S-reactive memory B cell frequency was significantly higher than that of HIV-1 Env and HCV E1E2 (Fig. 1D, 0.0088% and 0.0094%, respectively). We further confirmed the presence of pre-existing SARS-CoV-2 S-reactive humoral immunity by observing significantly higher plasma Ig levels against SARS-CoV-2 S compared to SARS-CoV S, MERS-CoV S and HIV-1 Env-specific Igs (Fig. S2E-F). The majority of the SARS-CoV-2 S-binding classical memory B cells were IgG+. In contrast, memory B cells reactive with the vaccine antigen TT were frequently IgM+, while H1N1 HA and RSV F reactive B cells were almost exclusively IgM-, probably reflecting recurrent antigenic stimulation (Fig. 1E, right panel; Fig. S2C). Similar results were obtained for the unswitched memory compartment, with SARS-CoV-2 having a higher S-reactive B cell frequency (0.042%) than that of HIV-1 Env (0.0083%) and HCV E1E2 (0.0018%), comparable to RSV F (0.034%), but lower than to H1N1 HA (0.076%) and TT (0.26%) (Fig. S2B). Thus, a significant portion of B cells in unexposed individuals is reactive with SARS-CoV-2 S. These B cells are found in multiple B cell compartments, suggesting that both *de novo* activation of naive B cells as well as reactivation of memory B cells elicited by a previous (HCoV) infection might play a role in the development of antibody responses after SARS-CoV-2 infection and/or vaccination. However, the majority of B cells that recognize SARS-CoV-2 are naive (82% of SARS-CoV-2 S-reactive B cells), whereas B cells against recurring infections and the TT vaccine predominantly have a memory phenotype, as one might expect (Fig. 1E, left panel; Fig. S2A, D).

To further evaluate the pre-existing SARS-CoV-2 reactive memory B cells, peripheral blood mononuclear cells (PBMCs) were stimulated with resiquimod (TLR7/8 agonist), IL-2 and IL-21 in order to promote the differentiation of memory B cells into plasmablasts and subsequently assess SARS-CoV-2 S-reactive Ig secretion as well as to a set of control antigens (Fig. 1F) (Franke et al. 2020). Stimulation of memory B cells resulted in the secretion of high antibody levels against viral proteins from previously encountered common seasonal infections (H1N1 HA, RSV F, endemic HCoVs), while low antibody levels could be detected against SARS-CoV S, MERS-CoV S, HIV-1 Env and HCV E1E2 (Fig. 1F). Stimulated memory B cells produced a significantly higher antibody titer against SARS-CoV-2 S than SARS-CoV S, HIV-1 Env and HCV E1E2, in line with memory B cell frequencies as measured by flow cytometry (Fig. 1D). In eight out of ten supernatants from unexposed individuals, the secretion of specific antibodies against SARS-CoV-2 S was similar to that against endemic HCoVs HKU1-CoV S and OC43-CoV S (Fig. 1F-H). Taken together, while the majority of SARS-CoV-2 S-reactive B cells belong to the naive compartment, the memory B cell compartment of the majority of HDs has the capacity to produce SARS-CoV-2 S-reactive antibodies despite not having previously encountered SARS-CoV-2. One to two donors appear to show some response to HIV-1 Env and HCV E1E2 (Fig. 1G-H), while they were confirmed HIV-1 and HCV-naive, an observation that requires further study but may relate to rare cross-reactivity of virus-reactive antibodies (Williams, Han, and Haynes 2018).

### S-reactive B cells often carry a IGHV1-69/IGKV3-11-derived BCR and are frequently polyreactive

Next, we single-cell sorted SARS-CoV-2 S-reactive B cells from four unexposed donors (HD01-HD04). The majority of the sorted cells were IgD^+^/CD27^-^ and thus belonged to the naive B cell compartment, confirming our previous observations (Fig. 2A). We obtained 132 heavy chain (HC) with 101 paired light chain (LC) sequences from the SARS-CoV-2 S-double positive B cells (Table S2). Of the 132 HC sequences obtained, 101, 20 and 5 were derived from naive, unswitched and classical memory B cells, respectively, while we were unable to attribute six to a defined B cell subset. The BCRs of the SARS-CoV-2 S-reactive naive B cells were virtually identical to their assigned germlines, while BCRs from all classical memory and some unswitched memory S-reactive B cells showed higher mutation rates, consistent with maturation in response to previous antigen exposure (Fig. 2B). In three of the four donors, IGHV1-69 was the most dominant HC V gene used by SARS-CoV-2 S-reactive B cells, while it was the second most dominant in the fourth donor, accounting for 39% of all expressed V genes across the four donors (Fig. 2C, left panel), representing an enrichment of up to tenfold compared to the unselected germline repertoire, where IGHV1-69 is expressed in 4-7% of BCRs (DeKosky et al. 2016; Briney et al. 2019). In the LC sequences a more moderate fourfold enrichment was observed for the IGVK3 family, which made up 36% of the LCs compared to 8% in an unselected naive repertoire (Fig. 2C, right panel) (DeKosky et al. 2016). When studying HC/LC pairing, we observed that 13/101 (13%) of all recovered pairs from SARS-CoV-2 S-reactive B cells are IGHV1-69/IGKV3-11 (Fig. 2D). While informative, the numbers of BCRs obtained here were quite low. Therefore, the BCR repertoire of SARS-CoV-2 S-reactive B cells was studied more extensively in the next section.

**Figure 2.**
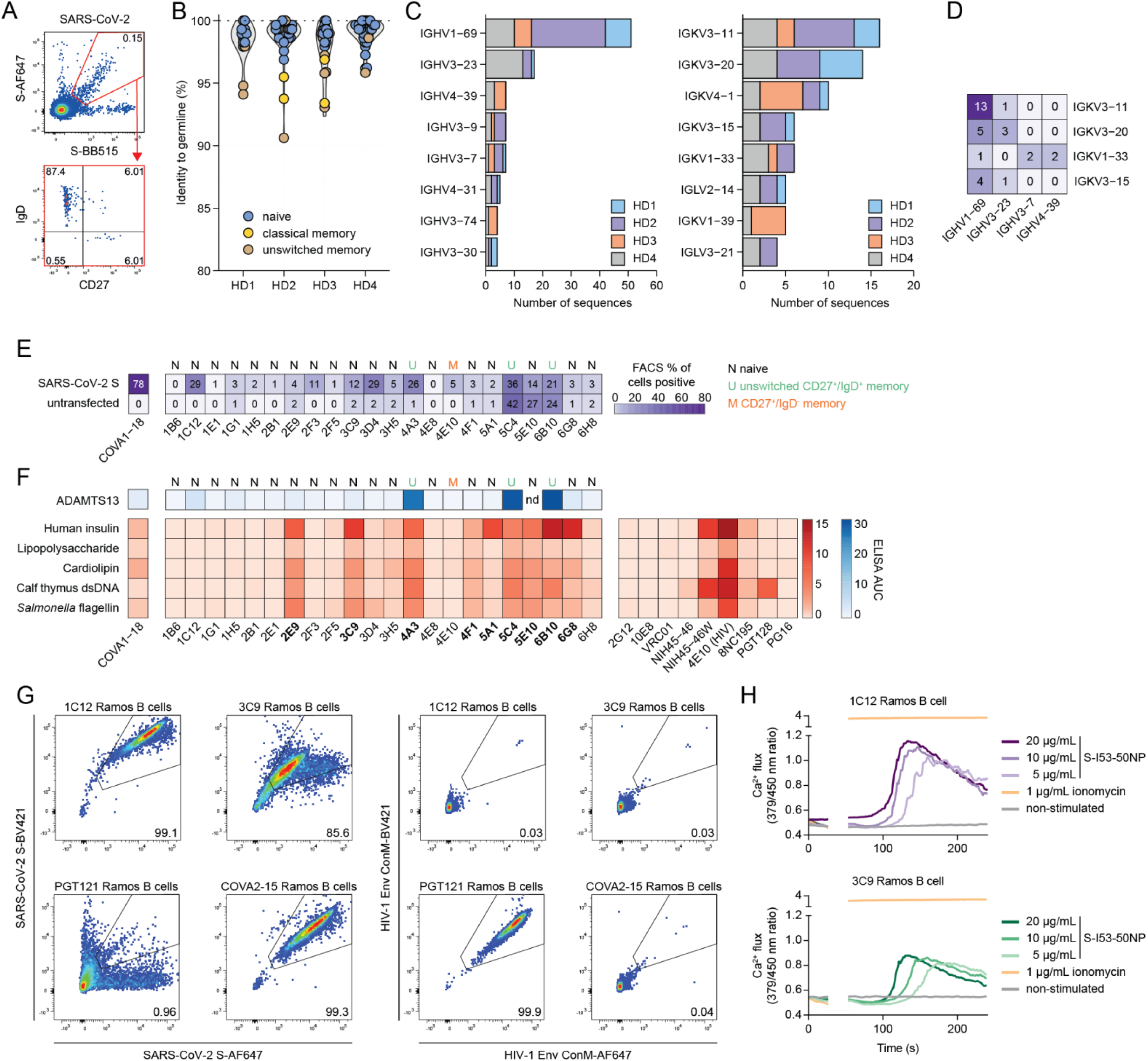
Genotypic and phenotypic characterization of SARS-CoV-2 S-reactive monoclonal antibodies from unexposed individuals. (**A**) Representative gating strategy of SARS-CoV-2 S-reactive B cells. Each dot represents an individual B cell. From antigen-specific B cells, the phenotype was determined (IgD^+^/CD27^-^, naive; IgD^+^/CD27^+^, unswitched IgD^+^ memory; IgD^-^/CD27^-^, CD27^-^ memory; IgD^-^/CD27^+^, classical memory) (lower panel). The numbers inside the boxes represent the frequency (%) of cells in a gate. (**B**) Dot plot overlaying a violin plot showing sequence identity (%) to IMGT-annotated germline heavy chain sequences for all isolated BCR heavy chains. Each color represents a B cell phenotype. HD, healthy donor. (**C**) Bar plot showing the number of sequences recovered for each immunoglobulin heavy chain V (IGHV, left panel) gene and immunoglobulin kappa/lambda light chain (IGKV/IGLV, right panel) gene. Colored bars represent different HDs. (**D**) Matrix showing the number of pairs with a certain IGHV (x-axis) and IGKV (y-axis). The numbers inside the boxes represent the number of pairs recovered for each pairing. (**E**) Matrix showing flow cytometric binding assay to SARS-CoV-2 S-transfected or untransfected HEK293T cells for each selected MAb and control MAb COVA1-18. Numbers and colors in the boxes represent the percentage of cells showing binding to a particular MAb. The phenotype as determined in (A) is shown on top of the matrix. (**F**) Matrix depicting area under the curve (AUC) as determined by a polyreactive enzyme-linked immunosorbent assay (ELISA) for each of the antigens shown on the left. The letters on top represent B cell phenotypes as determined in (A). Polyreactive MAbs are indicated in bold text. nd, not determined. (**G**) Antigen specificity of Ramos B cells designed to express a 1C12, 3C9, PGT121 or COVA2-15 BCR to SARS-CoV-2 S (left) or HIV-1 Env (right). The numbers inside the boxes represent the frequency (%) of cells in a gate. (**H**) Ramos B cell activation of 1C12 B cells (top panel) and 3C9 B cells (bottom panel) as measured by calcium (Ca^2+^) flux assay. A baseline without antigen was established between 0 and 30 seconds, after which the measurement was interrupted to add the antigen to the B cells (30-50 seconds). Ionomycin was used at 1 μg/mL as positive control.

The 101 HC/LC pairs were used to generate MAbs which were screened for binding to SARS-CoV-2 S by ELISA. 22 MAbs that showed significant binding in ELISA, defined as being three times above background, were selected for further evaluation. Of these 22 MAbs, 18 were from naive B cells, three from unswitched memory B cells and one from a classical memory cell. Three naive MAbs had the aforementioned IGHV1-69/IGKV3-11 pairing (MAbs 2F3, 3D4 and 5E10). The majority of the selected MAbs recognized cell surface-expressed full-length SARS-CoV-2 S (Fig. 2E and Fig. S3A), with varying efficiency. The binding was substantially lower compared to the control MAb COVA1-18 obtained from a SARS-CoV-2 convalescent individual (Brouwer et al. 2020), consistent with the fact that most of these MAbs were derived from naive B cells, and all were derived from B cells that had never encountered SARS-CoV-2. None of the 22 MAbs displayed any neutralizing activity in a pseudovirus neutralization assay (Fig. S3B). 5C4, 5E10 and 6B10 exhibited strong binding to untransfected cells (Fig. 2E and Fig. S3A), suggesting poly- and/or autoreactive properties. Therefore, the selected 22 MAbs were tested for polyreactivity against human insulin, bacterial lipopolysaccharide (LPS), bovine cardiolipin, calf thymus double-stranded DNA (dsDNA) and flagellin (Guthmiller et al. 2020) known polyreactive HIV-1 MAbs were included as controls. Nine MAbs showed some degree of polyreactivity, with the MAbs binding to untransfected HEK293T cells showing the highest degree of polyreactivity (Fig. 2F). In contrast, MAbs that showed a high degree of specificity in the cell-surface binding assay also showed little polyreactivity (e.g. 1C12). In addition, three of nine polyreactive MAbs were strongly cross-reactive against a disintegrin and metalloproteinase with a thrombospondin type 1 motif member 13 (ADAMTS13), an enzyme associated with COVID-19 thrombosis pathology (Fig. 2F) (Mancini et al. 2021; Doevelaar, Bachmann, Hölzer, Seibert, Rohn, Bauer, et al. 2021). All three were isolated from different HDs and originated from the unswitched memory compartment. Altogether, these results indicate that pre-existing anti-SARS-CoV-2 S-reactive B cells may have a high degree of polyreactivity, expansion of these polyreactive MAbs might play a role in thrombotic thrombocytopenia seen in COVID-19 disease and vaccination (Mancini et al. 2021; Doevelaar, Bachmann, Hölzer, Seibert, Rohn, Bauer, et al. 2021; Ruhe et al. 2021; de Bruijn et al. 2021).

To confirm that both polyreactive and non-polyreactive MAbs could be activated upon encounter with SARS-CoV-2 S as could occur during infection or vaccination, we engineered Ramos B cells to express a BCR that incorporates the variable regions of MAbs 1C12 and 3C9, both originally derived from naive S-reactive B cells. These MAbs are on opposite ends of the polyreactivity scale (1C12, low; 3C9, high) and show differential binding to S and thus represent diverse B cells that might encounter SARS-CoV-2 S *in vivo* (Fig. 2F). Both B cell lines recognized SARS-CoV-2 S (Fig. 2G) and were activated by I53-50 nanoparticules presenting 20 SARS-CoV-2 S and soluble SARS-CoV-2 S (Brouwer et al. 2021), irrespective of their polyreactive properties (Fig. 2H and Fig. S3C).

### IGHV1-69/IGKV3-11-derived BCRs are enriched in S-reactive naive B cells

To elaborate on the BCR signatures of SARS-CoV-2 S-reactive B cells observed above, we sorted additional S-reactive B cells starting from 40×10^6^ PBMCs of each donor (HD01-10) and performed single-cell sequencing using the 10X Genomics platform. We cross-referenced the variable regions with those of the ImMunoGeneTics (IMGT) database to obtain isotype, V(D)J gene usage and SHM, defined as mismatch to their assigned germline sequences (Lefranc et al. 2015). Next, we assigned each B cell to a naive or memory phenotype using k-means clustering based on the expression of surface markers IgD and CD27, defined with feature barcode antibodies, as well as the number of mismatches to its assigned germline V(D)J genes (Fig. 3A). Naive B cells (IgD^high^) were generally very close to their assigned germlines, whereas memory B cells (CD27^high^) exhibit higher levels of SHM and are generally IgD^low^ (Fig. 3A, left panel). The majority of all SARS-CoV-2 S-reactive B cells in HDs belong to the naive compartment (1755 naive vs. 210 memory), consistent with the results from our earlier flow cytometry data (Fig. 1). Similar numbers of naive and memory B cells were obtained for each HD, with the provision that HD05 yielded low B cell numbers overall (Fig. 3A, right panel). Many B cells in the memory cluster had class-switched, while nearly all B cells in the naive compartment only expressed IgD and/or IgM BCRs. Most of the IgM^+^ memory B cells are IgD^high^/CD27^high^, consistent with an unswitched memory phenotype (Fig. 3B).

**Figure 3.**
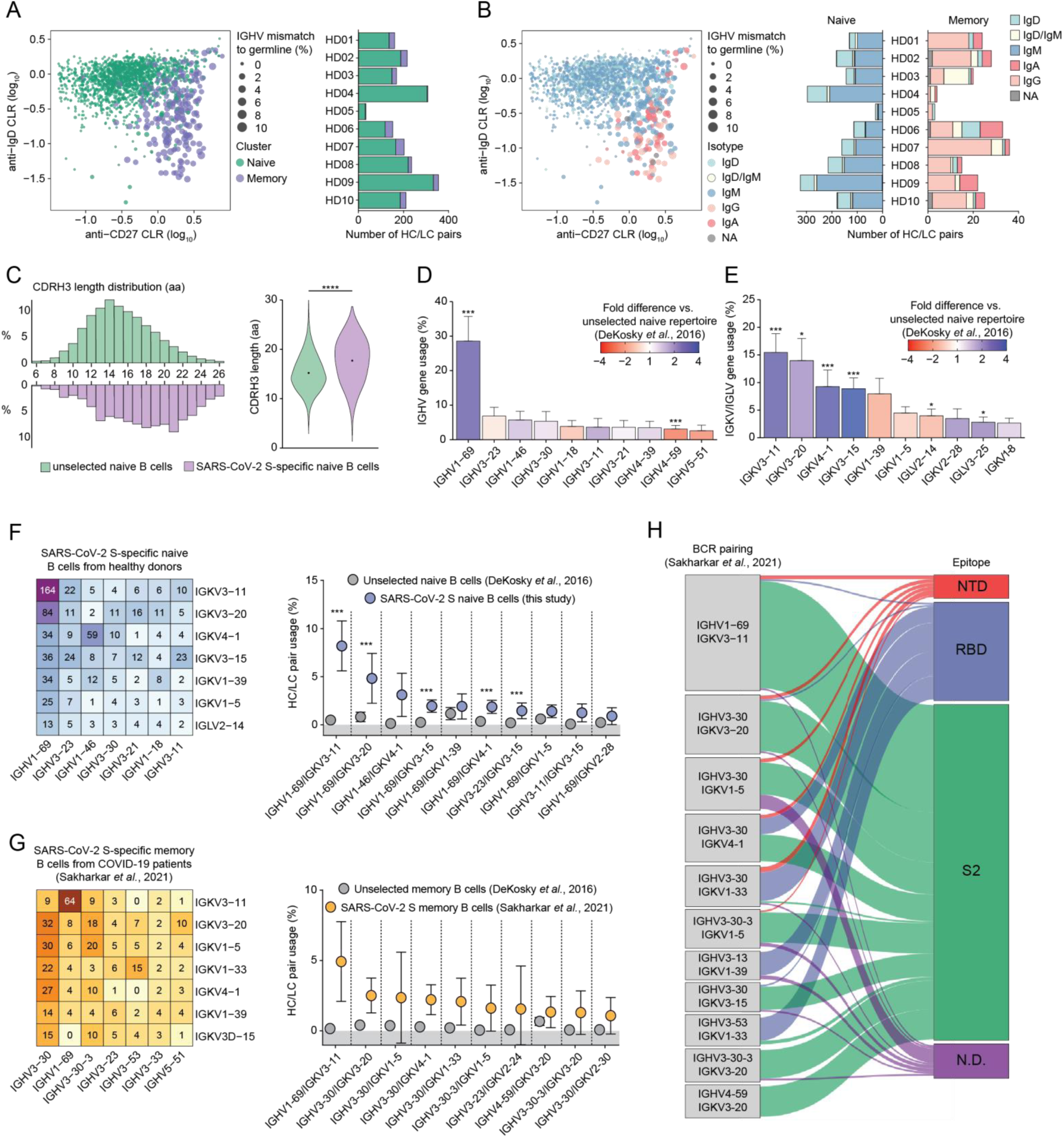
A public antibody class dominates the SARS-CoV-2 S-reactive B cell repertoire. (**A**) Jittered dot plot showing the phenotype of individual B cells and bar plot showing the number of recovered sequences per HD per cluster. k-means clustering was performed with three variables: anti-IgD centered log-ratio (CLR) normalized counts, anti-CD27 CLR normalized counts and number of total mutations in IGHV/IGKV/IGLV genes. Each dot represents an individual B cell, with the size of the dot corresponding to the percentage of mismatches to the IMGT-assigned germline sequence for IGHV genes. (**B**) Dot plot showing the isotype of individual B cells and bar plot showing the number of recovered sequences per HD per isotype, separated per cluster as determined in (A). NA, not applicable; if no isotype could be retrieved. (**C**) Heavy chain complementarity-determining region 3 (CDRH3) lengths in amino acids (aa) for naive SARS-CoV-2 S-reactive B cells from ten HDs (purple) and an unselected naive B cell repertoire (green). Significance was calculated with a Mann-Whitney U test. (DeKosky et al. 2016) (bottom panel). (**D-E**) Bar plots depicting the mean ± SEM IGHV (**D**) and IGKV/IGLV (**E**) gene usage (%) in naive SARS-CoV-2 S-reactive B cells from nine HDs. The colors indicate a positive (blue) or negative (red) fold difference over an unselected naive B cell repertoire (DeKosky et al. 2016). A non-parametric Kruskal-Wallis test was performed to compare the SARS-CoV-2 S-reactive naive repertoire to the unselected naive repertoire and is indicated on top of the bars. (**F**) Matrix showing the number of pairs with a certain IGHV (x-axis) and IGKV (y-axis) recovered from SARS-CoV-2 S-reactive naive B cells (left panel) and the frequency of observed BCR pairs compared to an unselected naive repertoire (right panel, mean ± SD) (DeKosky et al. 2016). The numbers inside the boxes represent the number of pairs recovered for each pairing. A non-parametric Kruskal-Wallis test was performed to compare the SARS-CoV-2 S-reactive naive repertoire to the unselected naive repertoire. (**G**) Matrix showing the number of pairs with a certain IGHV (x-axis) and IGKV (y-axis) recovered from SARS-CoV-2 S-reactive memory B cells from COVID-19 patients (Sakharkar et al. 2021) (left panel) and the frequency of observed BCR pairs compared to an unselected naïve repertoire (right panel, mean ± SD) (DeKosky et al. 2016). The numbers inside the boxes represent the number of pairs recovered for each pairing. No significance could be calculated since the unselected memory repertoire was n = 2. (**H**) Sankey diagram showing the most frequent BCR pairs in SARS-CoV-2 S-reactive memory B cells from COVID-19 patients and their epitopes from Sakharkar *et al*. (Sakharkar et al. 2021). N.D., not determined; *, *p* < 0.05; **, *p* < 0.01; ***, *p* < 0.001; ****, *p* < 0.0001).

We further studied the genetic characteristics of SARS-CoV-2 S-reactive naive B cells specifically, as only limited numbers of memory B cells were obtained. Heavy chain complementarity determining region 3 (CDRH3) length analysis revealed that the SARS-CoV-2 S-reactive naive B cell repertoire has on average longer CDRH3s than an unselected naive repertoire (Fig. 3C, Mann-Whitney U test, 17.7 vs 15.2 amino acids on average, respectively). This result is in line with previous findings on SARS-CoV-2 S-reactive memory B cells after infection (Briney et al. 2019; Brouwer et al. 2020; Barnes et al. 2020).

In accordance with the gene usage of isolated MAbs described above (Fig. 2C), S-reactive naive B cells showed a highly skewed HC gene usage in all HDs, with close to 30% of all sequenced HC genes belonging to the IGHV1-69 family (Fig. 3D), which is a significant enrichment compared to an unselected naive repertoire (3-fold, *p* = 0.009). IGHV1-46 and IGHV3-11 were also enriched, yet not significantly so (*p* > 0.05). In contrast, IGHV4-59 usage was significantly underrepresented (2.5-fold) compared to an unselected naive repertoire (Fig. 3D, *p* = 0.009). Four kappa light chain genes (i.e. IGKV3-11, IGKV3-20, IGKV4-1, IGKV3-15) were preferentially used compared to an unselected repertoire with IGKV3-11 representing over 15% of the BCR LC repertoire (Fig. 3E), consistent with our earlier observations (Fig. 2C). Indeed, usage of all three members of the IGKV3 Ig subfamily was significantly increased by at least 2-fold, as was IGKV4-1 (Fig. 3E, *p* = 0.009 for IGKV3-11, IGKV4-1 and IGKV3-15; *p* = 0.03 for IGKV3-15). SARS-CoV-2 S-reactive memory B cells of HDs do not show preferential gene usage compared to the unselected memory B cell repertoire, likely in part due to the low number of memory cells recovered (Fig. S4A-C). In summary, we observed substantial and significant increases in the usage of certain Ig genes, most notably in IGHV1-69, which is commonly involved in the humoral response to viral infections (Chen et al. 2019; Brouwer et al. 2020).

Next, we analyzed whether specific HC/LC pairings were preferred in SARS-CoV-2 S-reactive naive B cells in unexposed individuals. Indeed, we observed significant enrichment of multiple HC/LC pairings, in particular IGHV1-69/IGKV3-11, which was found in 164 out of 1755 total naive HC/LC pairs in S-reactive B cells (9.3%), corresponding to 8.5% on average for the nine donors included (Fig. 3F). These results match our earlier observations (Fig. 2D). Moreover, IGHV1-69 was used in seven out of the ten highest-frequency HC/LC pairings (IGHV1-69 paired with IGKV3-11, IGKV3-20, IGKV3-15, IGKV1-39, IGKV4-1, IGKV1-5, IGKV2-28) and four of these pairings were significantly overrepresented in SARS-CoV-2 S-reactive naive B cells compared to unselected naive B cells (IGHV1-69 paired with IGKV3-11, IGKV3-20, IGKV3-15, IGKV4-1, *p* = 0.009, Fig. 3F).

### IGHV1-69/IGKV3-11-derived BCRs are overrepresented in S-reactive memory B cells following SARS-CoV-2 infection

To contextualize these enriched naive clonotypes and verify whether they were relevant to the response following SARS-CoV-2 infection, we analyzed a recently published data set containing 1213 paired BCR sequences from SARS-CoV-2 S-reactive memory B cells isolated one to five months after SARS-CoV-2 infection (Fig. 3G) (Sakharkar et al. 2021). Strikingly, IGHV1-69/IGKV3-11 was the most frequent pair, representing 5.6% of the SARS-CoV-2 S-reactive memory B cell pool and showing ∼37-fold enrichment compared to the unselected memory repertoire. Most of the other pairings in SARS-CoV-2 S-reactive memory B cells involved IGHV3-30 (Fig. 3G). These observations highlight that IGHV1-69/IGKV3-11 is not only the predominant pair in the naive SARS-CoV-2 S-reactive B cell repertoire, but also expands and persists after SARS-CoV-2 infection.

Sakharkar and colleagues determined the epitope of these SARS-CoV-2 S-reactive memory B cells (Sakharkar et al. 2021), which allowed us to interrogate epitope specificities of certain enriched IGHV/IGKV pairings. From this dataset, most of the expressed MAbs targeted the S2 domain, while the RBD is targeted by the well-documented involvement of IGHV3-53 (Fig. 3H) (Barnes et al. 2020; Wu et al. 2020). The enriched IGHV1-69/IGKV3-11 pairing in SARS-CoV-2 S-reactive memory B cells is almost exclusively targeting the S2 domain, with only 4/64 (6%) targeted other epitopes on the S glycoprotein, and accounted for 9.3% of all S2-directed MAbs (Fig. 3H). These results suggest that a large pool of restricted IGHV1-69/IGKV3-11 naive B cells recognizes the S2 domain of SARS-CoV-2 S and that this B cell population is subject to expansion upon infection with SARS-CoV-2.

### IGHV1-69/IGKV3-11 B cells isolated from COVID-19 patients target an S2 epitope exposed on aberrant forms of S and can exert effector functions

To characterize the epitopes of the IGHV1-69/IGKV3-11 clonotype, we studied six non-clonal IGHV1-69/IGKV3-11 MAbs isolated from two COVID-19 patients in an earlier study (Brouwer et al. 2020), which all showed high identity to germline V genes and bear an average-length CDRH3 (Fig. 4A). Out of the six IGHV1-69/IGKV3-11 MAbs only COVA2-17 had the ability to neutralize SARS-CoV-2 pseudovirus. COVA2-17 stood out in this selection as it used a different D gene compared to the other five (IGHD2-15 versus IGHD3-22) and had a shorter CDRH3 loop (13 amino acids versus 15 or 16) (Fig. 4A). Previous studies revealed that COVA2-17 recognized the NTD, while the epitopes of the other five MAbs were not yet identified (Rosa et al. 2021). ELISA experiments revealed that the other MAbs bound to SARS-CoV-2 S, and all but COVA1-04 bound soluble S2 (Fig. S5A). Our observation that four out of six IGHV1-69/IGKV3-11 MAbs recognized S2, one recognized NTD, and one unresolved, is in accordance with the findings from Sakharhar *et al*. (Fig. 3H).

**Figure 4.**
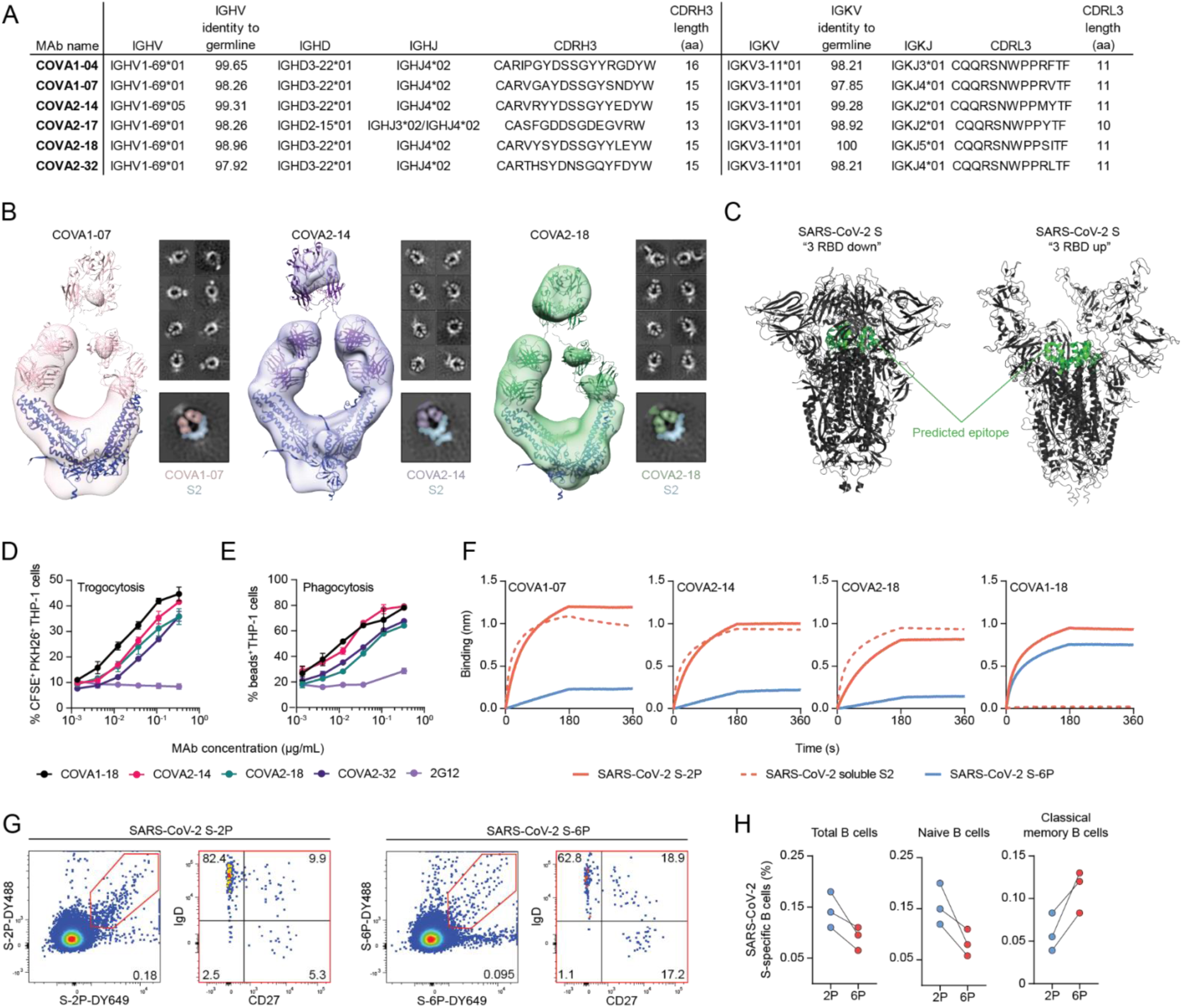
A public antibody class recognizes lesser stabilized epitopes on the S2 domain of SARS-CoV-2 S. (**A**) Characteristics of IGHV1-69/IGKV3-11 MAbs isolated in a previous study (Brouwer et al. 2020). (**B**) 3D reconstructions of COVA1-07 (pink), COVA2-14 (purple), and COVA2-18 (green) complexed with soluble S2. The models for IgG (PDB 1HZH) and the S2 domain (generated from PDB 7JJI) are docked into each density map. Individual representative 2D class averages of COVA MAbs in complex with S2 are shown to the right of each 3D reconstruction. (**C**) The predicted epitope (green) of COVA1-07, COVA2-14 and COVA2-18 MAbs on S2 in the context of SARS-CoV-2 S with 3 RBDs in the down conformation (PDB 7JJI, left) as well as 3 RBDs in the up conformation (PDB 7CAK, right). (**D-E**) Antibody-dependent cellular trogocytosis (**D**) and antibody-dependent cellular phagocytosis (**E**) assays for COVA MAbs and control MAb 2G12. (**F**) Biolayer interferometry (BLI) sensorgrams of COVA1-07, COVA2-14, COVA2-18 and COVA1-18 binding to SARS-CoV-2 S-2P (orange), SARS-CoV-2 soluble S2 (dashed orange) and SARS-CoV-2 S-6P (blue). (**G**) Representative gating strategy of SARS-CoV-2 S2P- and S6P-specific B cells. Each dot represents an individual B cell. From antigen-specific B cells, the phenotype was determined (IgD^+^/CD27^-^, naive; IgD^+^/CD27^+^, unswitched IgD^+^ memory; IgD^-^/CD27^-^, CD27^-^ memory; IgD^-^/CD27^+^, classical memory). The numbers inside the plots represent the frequency (%) of cells in a gate. (**H**) Connected dot plots showing the frequency of total (left panel), naive (middle panel) and memory (right panel) of SARS-CoV-2 S-2P- or S-6P-specific B cells (%) in three HDs.

Three of four S2-targeting MAbs (COVA1-07, COVA2-14 and COVA2-18) were then complexed as whole IgG molecule with soluble S2 and imaged by single-particle negative-stain electron microscopy (NS-EM, Fig. 4B). The low-resolution 3D reconstructions revealed an unusual topology in which the two Fab arms bound individual protomers of S2 (Fig. 4B and Fig. S5B). Due to the flexibility of soluble S2, no density was obtained for the third protomer of the trimeric S2 construct. Strikingly, all three IGHV1-69/IGKV3-11 MAbs, derived from different clonal lineages and from different donors engaged the soluble S2 in a highly similar way. The epitope is buried on the prefusion, “three RBD down” S trimer, which is consistent with the inability of these MAbs to neutralize SARS-CoV-2 (Fig. 4C). However, this epitope might be more accessible in rare conformations of S where two or three RBDs are in the “up” conformation (Fig. 4C).

We assessed whether the IGHV1-69/IGKV3-11-derived MAbs COVA1-07, COVA2-14, COVA2-18 and COVA2-32 recognized full-length S expressed on the surface of cells. All four S2 MAbs engaged full-length S and the binding was similar to that of the neutralizing RBD-specific COVA1-16 and COVA1-18 MAbs (Fig. S6A). These data are in line with observations that wild-type full-length S on viral/cell membranes is unstable and deteriorates into open conformations and/or S2 stumps that are in the post-fusion conformation (Ke et al. 2020). We also evaluated binding to full-length, cell surface-expressed S of globally circulating SARS-CoV-2 variants-of-concern (VOCs) alpha, beta, gamma and delta (B.1.1.7, B.1.351, P.1 and B.1.617.2, respectively) as well as endemic HCoVs (NL63, OC43, HKU1, 229E), epidemic HCoVs (SARS and MERS) and bat CoVs (SHC014, WIV-1) (Fig. S6A). The four S2 MAbs bound efficiently to the VOCs, SARS-CoV and bat CoV SHC014, but not to any of the other HCoVs tested, indicating that the epitope targeted by these IGHV1-69/IGKV3-11-derived MAbs is conserved across sarbecoviruses, but not beyond the sarbecovirus subgenus (Fig. S6A). Since these S2 MAbs were able to engage cell-surface expressed S protein we investigated their ability to partake in antibody-dependent effector mechanisms. IGHV1-69/IGKV3-11 MAbs COVA2-14, COVA2-18 and COVA2-32 efficiently triggered antibody-dependent cellular trogocytosis by the monocytic cell line THP-1, as shown by the extraction of membranous filaments from S-expressing cells (Fig. 4D and Fig. S6B). Similar results were obtained with the RBD-targeting neutralizing MAb COVA1-18, while the HIV-1 specific MAb 2G12 was unable to do so. COVA2-14, COVA2-18 and COVA2-32 also triggered antibody-dependent cellular phagocytosis as shown by the uptake of S-coated beads (Fig. 4E and Fig. S6C).

Some SARS-CoV-2 vaccines currently in use, including the AZD1222 vaccine from Oxford/AstraZeneca, encode for S-WT and are expected to expose aberrant forms of S that expose the IGHV1-69/IGKV3-11-targeted S2 epitope. Other vaccines, including mRNA-1273/Spikevax (Moderna), BNT162b2/Comirnaty (Pfizer/BioNTech), Ad26.CoV2.S (Janssen), and NVX-CoV2372 (Novavax) use a pre-fusion stabilized form of S that includes two proline substitutions (S-2P; (Sanders and Moore 2021)). However, several studies have shown that even S-2P is unstable and can deteriorate into aberrant conformations, albeit less efficiently so than S-WT (Hsieh et al. 2020; Juraszek et al. 2021). Indeed, we found that COVA1-07, COVA2-14 and COVA2-17 bound efficiently to S-2P as measured by bio-layer interferometry (BLI). In fact, the binding kinetics to S-2P were similar to those of S2, revealing that the non-neutralizing S2 epitope is well exposed on S-2P. Next, we tested a more stabilized S trimer that incorporates four extra prolines compared to S-2P, amounting to six extra prolines compared to S-WT (S-6P or HexaPro S (Hsieh et al. 2020)). S-6P is more resistant to heat/freeze cycles while its increased stabilization did not impact the conformation of S2 (Hsieh et al. 2020). BLI experiments revealed a large decrease of the binding of the S2 MAbs COVA1-07, COVA2-14 and COVA2-18 to S-6P compared to S-2P while RBD-targeting NAb COVA1-18 bound equally well to both proteins (Fig. 4F), indicating that IGHV1-69/IGKV3-11 MAbs might indeed target open and/or aberrant conformations of S2 that are more frequently present on S-2P compared to S-6P.

To better understand the population of B cells that target open or aberrant conformations of SARS-CoV-2 S in unexposed individuals, we compared the reactivity of B cells of three HDs to S-2P and S-6P by flow cytometry (Fig. 4G). Overall, the frequency of S-2P-reactive B cells was 1.6-fold higher than that of S-6P-reactive cells (Fig. 4G and Fig. 4H left panel). Strikingly, the loss of reactivity toward S-6P is explained by the lower frequency of naive and unswitched memory B cells reactive to S-6P compared to S-2P (∼2-fold) (Fig. 4H middle panel and Fig. S6D, E), whereas the opposite was true for the classical memory B cell population (Fig. 4H right panel, and Fig. S6D). This implies that naive B cells and unswitched memory B cells but not classical memory B cells often target open or aberrant S glycoprotein conformations in SARS-CoV-2-naive individuals. Moreover, we observed that MAbs derived from S-reactive B cells isolated from these unexposed individuals show a significantly reduced reactivity with S-6P compared to S-2P, confirming the propensity of naive and unswitched memory B cells to target open and/or aberrant conformations of S (Fig. S6F). Taken together, these results show that in unexposed individuals, a major part of the SARS-CoV-2 S-reactive B cell repertoire targets non-neutralizing epitopes on S2 exposed on aberrant forms of SARS-CoV-2 S which can trigger antibody responses upon natural infection and vaccination.

## Discussion

In this study, we aimed to elevate our understanding of the origin and development of the humoral response against SARS-CoV-2 S following infection or vaccination. To this end, we characterized the S-reactive baseline B cell repertoire prior to SARS-CoV-2 exposure. We observed that an unexpectedly large proportion of B cells is reactive with SARS-CoV-2 S. Recognition in unexposed individuals is predominantly mediated by the naive B cell compartment, but substantial numbers of classical and in particular unswitched SARS-CoV-2 S-reactive B cells are also present. The naive SARS-CoV-2 S-reactive B cells show a significant and meaningful enrichment in their BCR genes, with a public IGHV1-69/IGKV3-11 pairing present in 9.3% of S-reactive naive B cells. This enriched genotype was observed after vaccination and after infection, where it increased in frequency over time (Dugan et al. 2021; Sakharkar et al. 2021) as well as in a database attempting to collect all available anti-S MAb sequences (CoVAbDab), suggesting that the vast majority of patients developed this memory B cell subset (Dugan et al. 2021; Raybould et al. 2020). The molecular characterization of IGHV1-69/IGKV3-11 derived MAbs, which do not have any neutralizing capacity, revealed that this class of MAbs targets an epitope on the apex of the S2 domain, which is exposed on open and/or aberrant conformations of the S glycoprotein. While it is currently unclear why specifically this IGHV1-69/IGKV3-11 pairing targets this epitope, IGHV1-69 encodes a hydrophobic CDRH2 which has been shown to be involved in recognition of viral glycoproteins (Chen et al. 2019).

We also identified a rare pre-existing memory B cell pool able to recognize SARS-CoV-2 S in unexposed individuals. Because of the scarcity of this population, the derived BCR sequence data set was too restricted to be linked with memory B cell clonotypes identified in COVID-19 patients. The detection of this pre-existing memory population is in line with previous studies showing a boosting of endemic HCoV S-reactive antibodies in sera following SARS-CoV-2 infection (Grobben et al. 2021) and the isolation of HCoV cross-reactive antibodies from memory B cells showing a high degree of maturation (Sakharkar et al. 2021; K.-Y. A. Huang et al. 2021; Amanat, Thapa, et al. 2021). Molecular characterization of the 210 paired BCR sequences from SARS-CoV-2 S-reactive memory B cells reported in our study could help identify additional cross-reactive epitopes. In addition, we report heterogeneity of this pre-existing memory B cell pool, with some donors showing only a limited memory B cell response toward SARS-CoV-2 S. A subset of SARS-CoV-2 S-reactive pre-existing memory B cells has been shown to cross-react with commensal microbiota antigens, which may at least partially explain the origin of this population (Liu et al. 2021). Further studies should assess the role of these pre-existing memory B cells and their implications for disease outcome.

The immunodominance of the IGHV1-69/IGKV3-11 class of non-NAbs recognizing a conserved open and/or aberrant S conformation in SARS-CoV-2 infection raises questions about their role in COVID-19 outcome, and whether we should aim to promote or prevent their generation during vaccination. The open and/or aberrant conformations of S that are targeted by these non-NAbs, are present on cells expressing membrane-associated S protein and are therefore likely available on infected cells as well (Letarov, Babenko, and Kulikov 2021). Consistent with this, these non-NAbs can mediate antibody-dependent effector mechanisms and thereby may play a role in the clearance of SARS-CoV-2. Indeed, recent studies highlighted the contribution of Fc effector function in the protection, clearance or limitation of symptoms in COVID-19, in which one of the studied Abs was a non-neutralizing targeting S2 (Shiakolas et al. 2021; Beaudoin-Bussières et al. 2021). Moreover, it has been shown that Fc effector functions are a key determinant of protection after vaccination in a macaque model (Gorman et al. 2021). We report that the S2-targeting IGHV1-69/IGKV3-11 MAbs elicit effector functions and that they cross-react with multiple distinct S glycoproteins, including variants of concern and SARS-CoV-like viruses isolated from bats, but not endemic HCoVs. Overall, such antibody responses could help to mount a quicker adaptive immune response against new variants and potential *de novo* SARS-CoV-like outbreaks.

In contrast, exposure of immunodominant non-NAb epitopes might disfavor the induction of NAbs, which currently represent the most convincing correlate of protection for SARS-CoV-2 vaccines (Corbett et al. 2021; Earle et al. 2021; Feng et al. 2021). Thus, an argument can be made to engineer away non-NAb epitopes such as the one described here, in order to favor the induction of NAbs. Indeed, stabilization of viral glycoproteins in the native prefusion conformation has brought substantial improvements in the induction of NAb responses against the respective viruses, in particular RSV and HIV-1 (de Taeye et al. 2015; McLellan et al. 2013). Similarly, several studies have shown that stabilization of S with two proline substitutions (S-2P) promotes stronger NAb responses and/or protection following immunization in animal models (Amanat, Strohmeier, et al. 2021; Yu et al. 2020; Klasse, Nixon, and Moore 2021). As a result, S-2P was incorporated in the mRNA-1273/Spikevax (Moderna), BNT162b2/Comirnaty (Pfizer/BioNTech), Ad26.CoV2.S (Janssen), and NVX-CoV2372 (Novavax) vaccines. However, even S-2P is not entirely stable (Juraszek et al. 2021) and can expose non-NAb epitopes on S2. We show that a substantial fraction of naive B cells can still engage S-2P whereas the binding of non-NAbs of the IGHV1-69/IGKV3-11 class to S is abrogated when using the further stabilized S-6P. Ongoing vaccine trials with S-6P should shed light on whether preventing the induction of IGHV1-69/IGKV3-11 non-NAbs benefits the protective capacity of vaccines.

Finally, S2-targeting non-NAbs could participate in the physiopathology of COVID-19. Indeed, it has been shown that chronic (over)activation of the immune system can lead to the development of autoimmune disease through the expansion of auto- and/or polyreactive clones (Wong et al. 2021). In fact, this phenomenon has been observed in SARS-CoV and SARS-CoV-2 infection (A. T. Huang et al. 2020), with auto-antibodies to self-antigens such as clotting factor ADAMTS13 contributing to thrombotic thrombocytopenia (Doevelaar, Bachmann, Hölzer, Seibert, Rohn, Witzke, et al. 2021). We show that MAbs derived from SARS-CoV-2 S-reactive naive and unswitched memory B cells show polyreactive properties and in some cases bind to ADAMTS13, suggesting a potential role of these pre-existing S-reactive B cells in clotting events during SARS-CoV-2 infection. Similar thrombotic events, though more sporadic, have also been observed after administration of SARS-CoV-2 vaccines (de Bruijn et al. 2021; Al-Ahmad, Al-Rasheed, and Shalaby 2021; Maayan et al. 2021; Sissa et al. 2021), urging further research into ADAMTS13-reactive B cell characteristics in order to understand and potentially treat clotting events related to SARS-CoV-2 infection and vaccination.

In conclusion, we characterized the baseline SARS-CoV-2 S-reactive B cell repertoire. We found that a public antibody class using IGHV1-69 and IGKV3-11 very commonly recognizes a non-neutralizing epitope on SARS-CoV-2 S2 and that this antibody class is selected after SARS-CoV-2 infection and vaccination. These findings have implications for COVID-19 physiopathology and may help guide further improvements in SARS-CoV-2 vaccine design.

## Limitations of the study

One limitation of this study is that the novel combinatorial staining approach used to stain human B cells does not allow for the study of cross-reactive B cells. Therefore, although comparison to other HCoVs would have been useful, we did not incorporate viral proteins from viruses with high sequence identity to SARS-CoV-2 S. Moreover, although the aim was to also establish an expression profile of naive B cells using single-cell sequencing, the number of reads from transcriptionally inactive naive B cells was insufficient to include results on gene expression profiles. In addition, SARS-CoV-2 S-reactive B cells are relatively scarce in peripheral blood. Thus, we did not recover sufficient numbers of BCR sequences from S-reactive memory B cells to be able to extract a signature as we did for naive B cells. The present study focuses on characterizing SARS-CoV-2 S-reactive naive human B cells bearing an IGHV1-69/IGKV3-11 paired BCR. Though we aimed to determine their epitope on S, the polyreactivity and other physicochemical properties of these naive MAbs have made any epitope determination unfeasible. Through using a set of memory MAbs isolated from infected patients with identical genetic signatures, we aimed to resolve the epitope of these MAbs by proxy; however, future studies aimed at resolving the exact Ab-epitope interface may be able to elucidate why the IGHV1-69/IGKV3-11 is predominantly targeted to this epitope.

## Supporting information

Table S1 - FACS panels

Table S2 - MAbs properties

## Acknowledgements

We thank Single Cell Discoveries (Utrecht, the Netherlands) for help with single-cell RNA sequencing. We thank Drs. Li Wu and Vineet N. Kewal Ramani from the NIH AIDS Reagent Program for kindly providing Ramos B cells and Dr. Andrew McGuire for kindly sharing the pRRL.EuB29 lentiviral vector that was used to transduce Ramos B cells. We thank Prof. S Marieke van Ham and Dr. George Elias (Sanquin, the Netherlands) as well as Dr. Juan J. Garcia Vallejo (Amsterdam UMC, the Netherlands) for their help in developing the combinatorial probe staining strategy. Characterization of the BCR repertoire using RESEDA was carried out on the Dutch national e-infrastructure with the support of SURF Cooperative.

## Author contributions

Conceptualization: MC, TGC, RWS, MJvG

Funding acquisition: MC, ABW, AHCvK, RWS, MJvG

Investigation: MC, TGC, MdG, JH, DG, GK, BDCvS, AJ, AIS, MG, PJMB, KvdS, YA, JC-P, JLS, WO, IB, JAB, MP, TPLB, JLT, JC, ICM, SWdT

Methodology: MC, TGC, MdG, JH, BDCvS, AJ, AHCvK, PDM, RWS, MJvG

Project administration: MC, TGC, MJvG Resources: AA, MB, GJdB

Supervision: MC, TGC, ABW, KS, AHCvK, PDM, RWS, MJvG

Writing – original draft: MC, TGC, RWS, MJvG

Writing – review & editing: all authors

## Data and materials availability

All data is readily available in the main text and supplementary materials. NS-EM reconstructions will be deposited to the Electron Microscopy Data Bank under deposition IDs D_1000260663-1000260666. All reasonable requests for code and materials used in this study should be directed to and will be fulfilled under an MTA by Prof. Rogier W Sanders and Dr. Marit J van Gils.

## Funding statement

Netherlands Organization for Scientific Research (NWO) Vici grant (RWS)

Bill & Melinda Gates Foundation, Collaboration for AIDS Vaccine Discovery (CAVD) grant INV-002022 (RWS)

Amsterdam UMC AMC Fellowship (MJvG) Bill & Melinda Gates Foundation

COVID-19 Wave 2 mAbs grant INV-024617 (MJvG)

Fondation Dormeur, Vaduz (MJvG, RWS)

Netherlands Organization for Scientific Research (NWO) Veni grant (192.114) (MC)

## STAR Methods

### Sample collection

Blood was collected from healthy blood donors (n=10) by a Dutch blood bank (Sanquin, Amsterdam, The Netherlands) between March 2019 and February 2020, prior to the first official case of COVID-19 in The Netherlands. Plasma samples were collected after centrifugation and peripheral blood mononuclear cells (PBMCs) were isolated by density gradient centrifugation using Ficoll-Paque Plus. Isolated PBMCs were cryopreserved at -80°C for future usage.

### Protein design and purification

All soluble proteins, including SARS-CoV-2 S-2P and S-6P (Brouwer et al. 2020; Hsieh et al. 2020), S glycoproteins from variants of concern (Caniels et al. 2021) and endemic HCoVs (Grobben et al. 2021), influenza A hemagglutinin (H1N1pdm2009, A/Netherlands/602/2009, GenBank: CY039527.2) (Aartse et al. 2021), RSV prefusion stabilized F (DS-Cav1) (McLellan et al. 2013), hepatitis C virus E1E2 and HIV-1 ConM Env (Sliepen et al. 2019) constructs with avi-tag and/or hexahistidine (his)-tag and/or strep-tag were expressed and purified as previously described (Brouwer et al. 2020). After purification, avi-tagged proteins were biotinylated with a BirA500 biotin-ligase reaction kit according to the manufacturer’s instruction (Avidity). Tetanus toxoid was purchased from Creative Biolabs (Vcar-Lsx003) and aspecifically biotinylated using EZ-Link Sulfo-NHS-LC-Biotinylation Kit (ThermoFisher) according to the manufacturer’s instruction.

### Probe preparation and staining

Biotinylated protein antigens were individually multimerized with fluorescently labeled streptavidin (BB515, BD Biosciences; BUV615, BD Biosciences; AF647, Biolegend; BV421, Biolegend) as described previously (Brouwer et al. 2020). Briefly, biotinylated proteins and fluorescently labeled streptavidin were mixed at a 2:1 protein to fluorochrome molar ratio and incubated at 4°C for 1 h. Unbound streptavidin conjugates were quenched with 10uM biotin (Genecopoiea) for at least 10 minutes. Individual labelled proteins were then equimolarly mixed at a final concentration of 45.5 pM. 2-4×10^7^ previously frozen PBMC samples were first depleted for T cells using CD3 selection kit II (StemCell) according to the manufacturer’s instruction. Enriched B cells were then stained in Eppendorf tubes with 50-100 μL of antigen probe cocktail for 30 min at 4°C, subsequently washed with FACS buffer (PBS supplemented with 1 mM EDTA and 2% fetal calf) and stained with the Live/DEAD dye together with MAbs coupled with fluorophores for FACS (Table S1) for an additional 30 min at 4°C. Stained samples were washed twice and acquired on the BD LSRFortessa^TM^ for cell analysis and ARIA-SORP-II 4 lasers for cell sorting. Analysis was performed using DIVA and Flowjo 10 software (BD Biosciences).

### Combinatorial probe staining strategy

A combinatorial probe staining strategy was set up to perform simultaneous identification of multiple B cell specificities in a single sample, in the context of limited parameters/channel, which has been used previously with pMHC tetramers (Hadrup et al. 2009). Conventionally, antigen-specific B cells are detected by the binding of two different fluorochrome-coded to the same protein. This method of double probe staining has been commonly used to reveal the fine specificity of B cells and avoid artefact of non-specific B cells binding to the fluorophore itself. Contrastingly, this combinatorial probe staining strategy uses all possible combinations of two fluorophores to increase the number of specificities that can be detected. The number of different B-cell specificities that can be detected equals N(N-1)/2, where *N* is the number of different fluorescent labels. In our study, we were able to detect 6 different antigen-specificities using 4 distinct fluorophores. Probes were labelled and used as previously described, in the following manner: SARS-CoV-2 S (AF647, BV421), H1N1 HA (BUV615, BV421), RSV F (AF647, BUV615), HCV E1E2 (AF647, BB515), HIV-1 ConM Env (BB515, BUV615) and Tetanus toxoid (BB515, BV421). Probe staining was used in combination with LIVE/DEAD dye (ThermoFisher) and labeled antibodies as previously described and acquired by FACS. For the analysis, the lymphocyte population was first gated based on the morphology (FSC-A/SSC-A) and doublets were removed. Next, dead cells and remaining CD4^+^ cells to avoid artefact binding of HIV-1 probes were first excluded within a dump channel and live antigen-specific B cells were studied in the CD19^+^population. To remove potential cross-reactive B cells to streptavidin, each probe combination was first gated on cells double negative for the two other channels (Fig S1).

### B cell stimulation

Polyclonal stimulation was performed according to the published method from (Franke et al. 2020). Briefly, the freshly thawed PBMC were resuspended in RPMI 1640 (Gibco) supplemented with 10% fetal bovine serum, 100 U/mL penicillin/streptomycin, 2 mM L-Glutamine, 1 mM sodium pyruvate, 8 mM HEPES (RPMI10) (Life Technologies, Grand Island, NY, USA). R848 (MedChemExpress), recombinant human IL-2, and IL-21 (ImmunoTool) were used at a final concentration of 1 µg/mL, 100 units/mL and 50 ng/mL, respectively. PBMCs were cultured at 2 × 10^6^ cells/mL, in 24-well Greiner CELLSTAR suspension culture plates and were incubated at 37 °C, 5% CO2 for 9 days. Harvested supernatants were then used for subsequent experiments.

### Luminex assay

Luminex assays were performed as described previously (Grobben et al. 2021). In short, expressed glycoproteins were covalently coupled to Luminex Magplex beads using a ratio of 75 µg protein to 12.5 million beads for SARS-CoV-2 S and equimolarly for all other proteins. Next, 1:10.000-diluted plasma or undiluted stimulated B cell supernatant was added to the protein-coated beads overnight at 4°C. The following day, beads were washed with TBS/0.05% Tween using a magnetic separator and resuspended in 1,3 µg/mL goat anti-human IgG-PE and incubated for 2 hours at room temperature (Southern Biotech). After washing, the beads were resuspended in Magpix drive fluid (Luminex). The plates were read on a Magpix (Luminex). Specific binding was attributed to median fluorescence intensity (MFI) levels >3x above background.

### B cell sorting for MAb isolation

SARS-CoV-2 S biotinylated proteins were individually multimerized with fluorescently labeled AF647 and BV421 streptavidin and used for staining as previously described. Enriched B cells were stained with probes and antibodies conjugated to fluorophore and LIVE/DEAD dye (ThermoFisher) (Table S1). The lymphocyte population was first gated based on the morphology (FSC-A/SSC-A) and doublets were removed. Dead cells and non-B cells were first excluded within a dump channel (CD3^-^/CD14^-^/CD16^-^). Live B cells (CD19^+^) that were double positive for the SARS-CoV-2 S protein (AF647 and BV421) were single cell-sorted using index sorting into a 96-well plate containing a lysis buffer. The lysis buffer consisted of 20 U RNAse inhibitor (Invitrogen), first strand SuperScript III buffer (Invitrogen), 1.25 μl of 0.1 M DTT (Invitrogen), in a total volume of 20 μL. The plates with the sorted single cells were stored at -80°C for at least 1 h before performing reverse transcriptase (RT)-PCR.

### B cell receptor variable region amplification, cloning and MAb expression

The BCR heavy chain and light chain variable regions were amplified and cloned into an IgG1 expression vector using reverse transcriptase polymerase chain reaction (RT-PCR) as described previously (Brouwer et al. 2020). The V(D)J variable regions of antibodies were then cloned into human IgG1 expression vectors. The cloning was performed by mixing 2 μL of home-made Gibson enzyme mix containing 5X isothermal reaction buffer (0,5 g PEG-8000, 1M Tris/HCl at pH 7.5, 1M MgCl_2_, 1M DTT, 100 mM of each dNTP, 50 mM NAD and MQ), 1U/μL T5 exonuclease, 2U/μL Phusion polymerase and MQ, together with 1 uL of enzymatically digested expression vector (50ng/uL) and 1 uL of PCR product. This mixture was then transformed into chemically competent *Escherichia coli*. After DNA purification, the sequences were verified by Sanger sequencing. For small scale expression, adherent HEK293T cells cultured in Dulbecco’s Modified Eagle Medium (DMEM) supplemented with 10% fetal calf serum and a mixture of penicillin/streptomycin (100U/mL and 100 μg/mL, respectively) were transfected with purified DNA as described previously (Brouwer et al. 2020). Supernatants of transfected cells were harvested 48 h post-transfection.

### Preliminary ELISA screening of MAb supernatants

Preliminary enzyme-linked immunosorbent assays (ELISA) were conducted with clarified supernatants from HEK293T cells transfections. In order to maintain equimolar binding, SARS-CoV-2 RBD- and S-his-tagged proteins were diluted in casein to 1 μg/mL and 4.8 μg/mL, respectively. Antigens were immobilized for 2 h at RT on Ni-NTA 96-well plates. After three washes with TBS, binding of supernatants was allowed for 2 h at RT. Horseradish peroxidase (HRP)-labelled secondary antibody (goat anti-human IgG 1:3000) in casein was then incubated for 1 h at RT. After five washes with 1xTBS/0.05% Tween-20, 100 μl of freshly-made developing solution (1% 3,3’,5,5’-tetramethylbenzidine, 0.01% hydrogen peroxide, 0.1 M sodium acetate and 0.1 M citric acid) were added to the plates. The reaction was terminated by adding 50 ul of 0.8 M sulfuric acid to each well and optical density at 450 nm (OD450) was measured.

### Large-scale expression of monoclonal antibodies

MAbs that showed binding in the preliminary screening ELISA assays were selected and produced at a larger scale. Briefly, 250 mL suspensions of HEK293F cells were maintained in FreeStyle medium and transfected with 19.5 μg of the two HC and LC plasmids in a 1:1 ratio, together with 117 μL of PEImax (1 mg/ml). The produced MAbs were then harvested and purified after five days. For antibodies purification, cell suspensions were centrifuged for 30 min at 4000 rpm, and then filtered using 0.22 μm pore size SteriTop filters (Millipore), followed by a run of the supernatants over a 1 ml protein G beads column. 18 ml of elution buffer (0.1 M glycine pH 2.5) were used to elute the antibodies into 2 ml of neutralization buffer (1 M TRIS pH 8.7). By using 100kDa VivaSpin20 columns, the purified antibodies were then concentrated and buffer exchanged to PBS. Concentration was measured on a NanoDrop 2000 (ThermoFisher).

### Binding assays to cell surface expressed CoV-S by Flow cytometry

HEK293T cells were transfected with full length S plasmid DNA (SARS-CoV-2 WT and variants, other epidemic and endemic CoVs, and Bat-CoVs) using Lipofectamine2000 (Invitrogen). Briefly, 0.5×10^6^ cells/well were plated in a 6-well plate. After 24h, 4 μg DNA and 10 μL lipofectamine were mixed, incubated and added to each well. After 48 h, cells were harvested and pooled, and 5×10^4^ cells were incubated in RPMI with 50 μg/mL of purified SARS-CoV-2 S-reactive MAbs for 1 h at RT. Cells were subsequently washed twice with PBS and stained for 30 min on ice and in the dark, in 50 μL of FACS buffer containing 1:1000 diluted PE-conjugated goat anti-human IgG (Biolegend). Cells were then washed twice with FACS buffer, fixed with 2% PFA and subsequently analyzed on BD LSRFortessa^TM^.

### Polyreactivity ELISA

Polyreactivity ELISAs were performed as described elsewhere (Guthmiller et al. 2020). High-binding half-area 96-well plates (Costar) were coated with 2 μg/mL *Salmonella enterica* serovar *Typhimurium* flagellin (Invitrogen), 10 μg/mL calf thymus double-stranded DNA (ThermoFisher), 5 μg/mL human insulin (Sigma-Aldrich) and 10 μg/mL *Escherichia coli* lipopolysaccharide (Sigma-Aldrich) in PBS and stored at RT overnight. Separate plates were coated with 10 μg/mL bovine cardiolipin (Sigma-Aldrich) in 99% ethanol and allowed to air-dry overnight. The following day, plates were washed with demineralized water and blocked with PBS/0.05% Tween/1mM ethylenediaminetetraacetic acid (EDTA) for 1 h. MAbs were serially diluted five-fold (starting concentration 10 μg/mL) and binding to the plates was allowed for 2 h at RT. After washing, HRP-labelled secondary antibody (goat anti-human IgG 1:3000) in PBS was then incubated for 1 h at RT. Plates were washed and developed as described before. All experiments were performed in duplicate and are representative of two independent experiments.

### Generation of Ramos B cells expressing custom B cell receptors

The B cell specific expression plasmid was constructed by exchanging the gl2-1261 gene of the pRRL EuB29 gl2-1261 IgGTM.BCR.GFP.WPRE plasmid (McGuire et al. 2014) with the variable heavy and light chain genes of 1C12, 3C9, PGT121 and COVA2-15 using Gibson assembly (Integrated DNA Technologies). The production of lentivirus in HEK293T cells and the subsequent transduction was conducted as described elsewhere (ter Brake et al. 2006). In short, lentiviruses were produced by co-transfecting the expression plasmid with pMDL, pVSV-g and pRSV-Rev into HEK293T cells using lipofectamine 2000 (Invitrogen). Two days post transfection, IgM-negative Ramos B cells cultured in RPMI10 were transduced with filtered (0.45 μm) and concentrated (100 kDa molecular weight cutoff, GE Healthcare) HEK293T supernatant. Seven days post-transduction, BCR-expressing B cells were sorted on IgG and green fluorescent protein (GFP) double-positivity using a FACS Aria-II SORP (BD biosciences). B cells were then expanded and cultured indefinitely.

### Ramos cell binding assay

Antigen specificity of the generated 1C12, 3C9, PGT121 and COVA2-15 Ramos B cells to SARS-CoV-2 S and HIV-1 ConM Env was detected as previously described using labelled probes and flow cytometry. Briefly, SARS-CoV-2 S and ConM v7 Env proteins multimerized with fluorescently labeled AF647 and BV421 streptavidin were used to stain 5×10^5^cells together with live/DEAD dye, IgM-BV605, IgG PE-Cy7, and the antigen-probe cocktail (Table S1). Stained samples were subsequently washed twice with FACS buffer and acquired on the BD LSRFortessa^TM^for cell analysis. Analysis was performed using FlowJo v10.7. Ramos cells were first gated based on the morphology (FSC-A/SSC-A) and doublets were removed. Live cells were selected and subsequently gated on IgM-, GFP+ and IgG+. (Antigen-specific ramos cells were double positive for the SARS-CoV-2 S protein (AF647 and BV421) and ConM Env protein (AF647 and BV421)).

### Calcium flux assay

B cell activation experiments of Ramos B cells were performed as previously described (Brouwer et al. 2021; Sliepen et al. 2019). In short, 4×10^6^ cells/mL in RPMI10 were loaded with 1.5 μM of the calcium indicator Indo-1 (Invitrogen) for 30 min at 37°C, washed with Hank’s Balance Salt Solution supplemented with 2 mM CaCl_2_, followed by another incubation of 30 min at 37°C. Antigen-induced Ca^2+^ influx of B cells was monitored on a LSR Fortessa (BD Biosciences) by measuring the 379/450 nm emission ratio of Indo-1 fluorescence upon UV excitation. Following 30 s of baseline measurement, aliquots of 1×10^6^ cells/mL were then stimulated for 210 s at RT with either 20 μg/mL, 10 μg/mL or 5 μg/mL of SARS-CoV-2 S or the equimolar amount presented on I53-50NPs. Ionomycin (Invitrogen) was added to a final concentration of 1 μg/μL to determine the maximum Indo-1-fluorescence. Kinetic analyses were performed using FlowJo v10.7.

### Flow cytometry for 10X Genomics

Enriched B cells from 4×10^7^ PBMCs of 10 donor were stained with Human TruStain FcX Fc Blocking Reagent (BioLegend, 422302) for 10 min at 4 °C. For each donor a mix of Abs linked to feature barcodes, containing one specific hashtag barcode together with CD27 and IgD, was centrifuged at 14,000*g* at 4 °C for 10 min and supernatant harvested. Barcoded antibody mix, anti-CD19-AF700, Live/DEAD dye, and labelled SARS-CoV-2 S (AF647 and BV421) were added to the cells and stained for 30min at 4 °C. Cells were then washed twice and resuspended in FACS buffer. Live B cells positives for the SARS-CoV-2 S protein (AF647 and BV421) were bulk sorted, for a total of 10,700 cells, and used for downstream 10X Genomics analysis.

### 10X Genomics library construction

Single-cell embedding in gel beads-in-emulsion (GEMs) and generation of barcoded cDNAwas performed on a 10X Genomics Chromium Controller, following the 10X Genomics v1.1 single-cell V(D)J Next GEM with Feature Barcoding technology for cell surface protein chemistry. In short, single cell suspensions were kept at 4°C and processed within 2 h upon completion of single cell sorting. Cells were then filtered to prevent clumping and counted to assess cell integrity and concentration. Out of 10,000 cells that were sorted, approximately 8,000 cells were loaded and the resulting sequencing libraries were prepared following standard 10X Genomics protocols, generating a transcriptome, cell surface protein marker and a V(D)J library from each experiment. The cDNA libraries were paired-end sequenced on an Illumina NovaSeq S4 with a 2×150 bp Illumina kit.

### Computational analyses for single cell sequencing data

Demultiplexing of raw base call (BCL) files, alignment, read filtering, barcode and UMI counting of the gene expression and feature barcode (10 hashtag oligos (HTOs) to discriminate individual donors and two antibody-derived tags (ADTs) for IgD/CD27 phenotyping) libraries was performed with the 10X Genomics Cell Ranger analysis pipeline using ‘cellranger count’. Sequence assembly of the V(D)J library was performed using ‘cellranger vdj’. During sequencing, Read 1 was assigned 28 base pairs and was used for identification of the Illumina library barcodes, cell barcodes and UMIs. Read 2 was used to map to the human reference transcriptome (GRCh38, version 3.0.0) and the CellRanger human V(D)J reference (GRCh38, version 5.0), respectively. Filtering of empty barcodes was done following standard CellRanger procedure. Resulting filtered feature-barcode matrices were imported in R (version 4.0.3) using the Seurat package (version 4.0.2) (Hao et al. 2021). HTO and ADT UMI counts were normalized using centered log ratio (CLR) transformation. The Seurat function ‘MULTIseqDemux’ was used to demultiplex samples from the 10 HDs using the HTOs and identify singlets, doublets and negative cells (McGinnis et al. 2019).

RESEDA (https://bitbucket.org/barbera/reseda/) was used to further analyze the BCR repertoire. Sequences assembled by CellRanger were aligned to the IMGT gene database (Giudicelli, Chaume, and Lefranc 2005) with BWA (default settings) (Li and Durbin 2010). Variants were called with samtools mpileup (Li et al. 2009) and VarScan (Koboldt et al. 2009) with minimum coverage set to 1 read to avoid missing mutations from antibody parts covered by a single sequence read. Consequently, any differences with respect to the IMGT sequences were considered as SHM. The CDR3 sequences were determined by translating the nucleotide sequences to peptide sequences and searching for conserved motifs in the V and J genes. The information from RESEDA was merged with the results obtained with Seurat.

### Clustering and BCR repertoire analysis

In total, 1965 paired heavy/light chain BCRs were recovered, which could be assigned reliably to a single HD using the HTOs. K-means clustering with three determinants (IgD CLR/CD27 CLR/total number of mutations in HC/LC V region) was used to cluster B cells. The optimal number of clusters (k = 2) was determined using the average silhouette method and was used to assign a phenotype (memory/naive). IMGT-assigned CDRH3 lengths were compared to those of a large dataset of naive B cells available online (Briney et al. 2019). For comparison of the expression of variable regions, paired HC/LC BCRs from naive and memory B cells were separately compared to naive and memory B cells in previously published data sets, respectively (DeKosky et al. 2016, Sakharkar et al. 2021). Analysis and handling of datasets was performed using R version 4.0.3.

### Negative-stain electron microscopy

MAbs in molar excess were complexed with S2 for 30 min at RT. Immune complexes were diluted to ∼20 μg/mL and deposited on glow-discharged, carbon-coated 400 mesh copper grids (Electron Microscopy Sciences). Sample was blotted from the grid with filter paper followed by two successive additions of 2% w/v uranyl formate stain with blotting. Grids were imaged on a Tecnai T20 (FEI) electron microscope with a CMOS 4k camera (TVIPS) at 200 kV, 62,000x nominal magnification, and 1.77Å/pixel. Micrographs were collected using Leginon, particles were picked using Difference of Gaussians picker, and particles were cleaned through successive rounds of reference-free 2D classification in Relion 3.0 (Suloway et al. 2005; Voss et al. 2009; Lander et al. 2009; Zivanov et al. 2018). Particles were also processed in CryoSPARC2 and reconstructed using Ab Initio (Punjani et al. 2017). Figures were made in UCSF Chimera and Adobe Photoshop (Pettersen et al. 2004).

### Biolayer interferometry

Nickel–nitrilotriacetic acid (Ni-NTA) biosensors (ForteBio) were loaded with 20 μg/mL his-tagged S glycoproteins (S-2P and S-6P) or equimolar amounts of his-tagged soluble S2 in running buffer (PBS, 0.02% Tween 20, and 0.1% bovine serum albumin) for 300 s. Subsequently, the biosensors were washed in a well containing running buffer to remove excess protein, followed by transfer to a well containing MAb (50 μg/mL) in running buffer for 180 s to measure association. Next, the biosensors were moved to a well containing running buffer for 180 s to measure dissociation of the S–MAb complexes. All measurements were performed on an Octet K2 (ForteBio).

### Antibody-dependent cellular trogocytosis

HEK293F cells (Invitrogen) at a density of 1×10^6^cells/mL were transfected using SARS-CoV-2 S plasmid and PEImax (1 µg/µl) in a 3:1 ratio in OptiMEM. HEK293F cells were harvested 72 hours after transfection and their plasma membrane was stained with 10µM PKH26 (Sigma-Aldrich) dye in PBS, for 20 min (RT) with periodic mixing. Cells were washed twice with PBS and taken up in culture medium. THP-1 effector cells (ATCC) were stained intracellularly with 0.05µM carboxyfluorescein succinimidyl ester (CFSE, ThermoFisher) in PBS and incubate 20 min (RT) with periodic mixing. Cells were washed twice with PBS and taken up in culture medium.). PKH26 stained HEK293F cells were opsonized for 30 min at 37°C, with serial MAb dilutions. 2G12-IgG1, specific for HIV-1 gp120, was used as a negative control. After incubation, cells were washed and THP-1 cells were added to the HEK293F cells at a 2:1 effector:target ratio. Plates were centrifuged shortly to promote cell to cell contact and incubated 1 hour at 37°C. Afterwards, cells were washed and resuspended in PBS/2% FCS. Flow cytometry was used to measure the double positive, PKH26^+^CFSE^+^THP-1 cells. ADCT was calculated by the fraction of THP-1 cells that received membrane fragments from the HEK293F cells. To exclude measurement of antibody-independent trogocytosis, cells were gated on stained HEK293F and THP-1 cells in the absence of antibodies.

### Antibody-dependent cellular phagocytosis

Fluorescent neutravidin beads (Invitrogen) were incubated with biotinylated SARS-CoV-2 S-2P or RBD protein overnight at 4 °C. Beads were subsequently centrifuged shortly and washed twice with PBS/2% BSA to remove unbound antigen and block the remaining hydrophobic sites on the microspheres. The coated beads were resuspended in PBS/2% BSA and 0.1 μL of the original suspension was placed in every well of a V-bottom 96 well plate and incubated (2 h at 37°C) with serial MAb dilutions. 2G12-IgG1, specific for HIV-1 gp120, was used as a negative control. After incubation, plates were washed and 5×10^4^THP-1 effector cells (ATCC) were added to each well in a final volume of 100 μL of RPMI10. Subsequently, plates were centrifuged shortly to promote beads to cell contact before incubation (5 h at 37°C). After incubation, the cells were washed, resuspended in PBS/2% FCS and analyzed by flow cytometry.

### Data analysis and visualization

Data visualization and statistical analyses were performed in GraphPad Prism software (version 8.3), while sequence handling, analysis and visualization were performed in RStudio (version 1.3.1093, R 4.0.3). A non-parametric Wilcoxon signed-rank test was performed to assess statistical differences for paired samples, while a Mann-Whitney U test was performed for unpaired samples. A Mann-Whitney U test to compare CDRH3 lengths was performed in RStudio 1.3.1093 (R 4.0.3). All FACS data analyses were performed in FlowJo 10.8.0. Significance is denoted as ****, p < 0.0001; ***, p < 0.001; **, p < 0.01; *, p < 0.05; ns, not significant.

---

**Figure S1.**
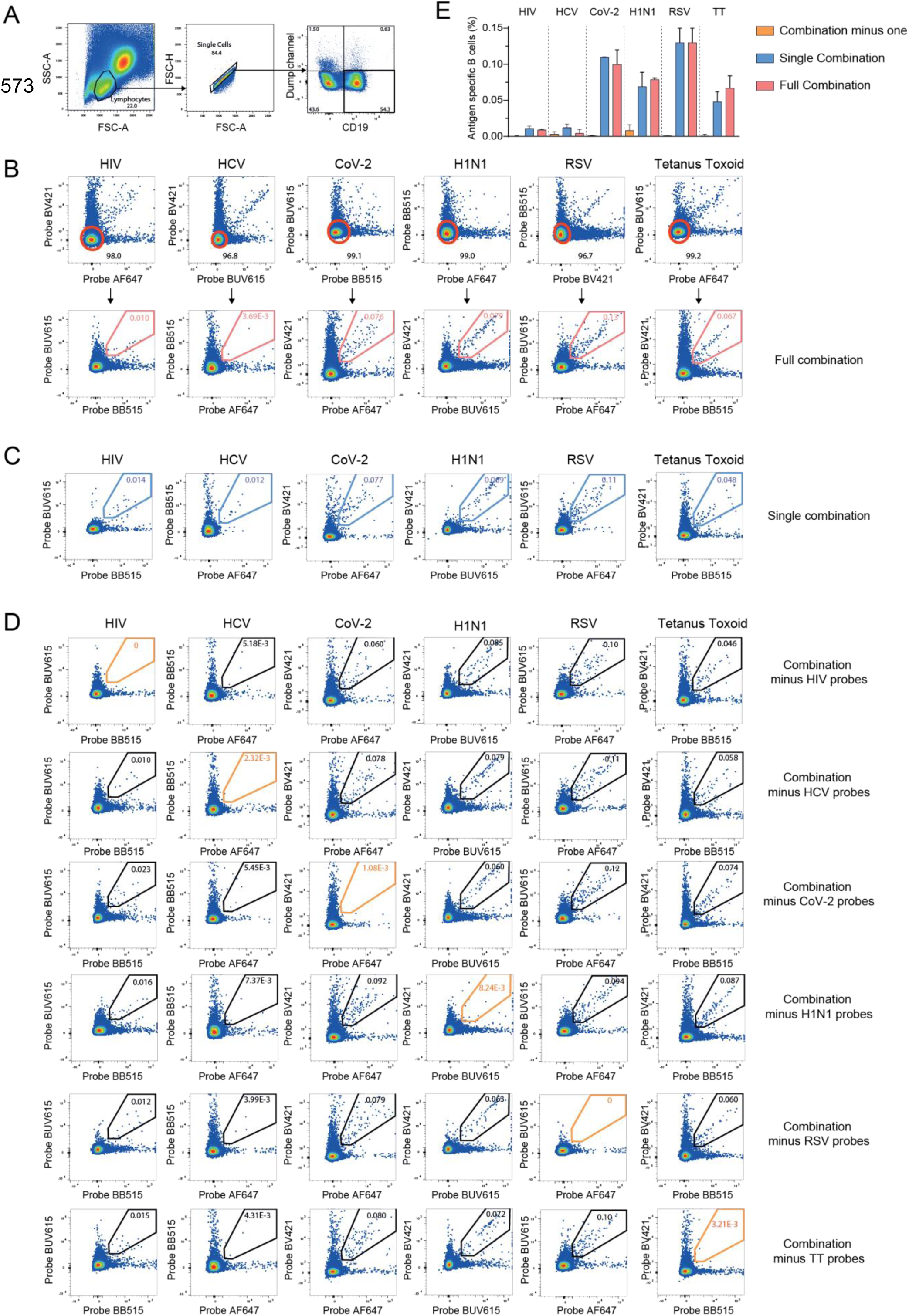
Combinatorial probe staining strategy validation. (**A**) Gating strategy to identify live B cells. **(B)** Combinatorial probe staining and gating strategy for the detection of multiple B cell specificities in a single PBMC sample (see method section). Top panel: Each dot represents a live B cell. To remove potential cross-reactive B cells to streptavidin, each probe combination was first gated on cells double negative for the two other channels. Bottom panel: Antigen-specific B cells are then detected as double positive for the binding of the same antigen multimerized with two different fluorochromes according to a matrix code (Fig. 1A), all combinations were performed in a single sample (full combination). (**C**) Single combination: each single combination of antigen multimerized with two different fluorochromes was performed in independent samples. (**D**) Combination minus one: All combinations of antigen multimerized with two different fluorochromes minus one were performed in a single sample, removal of each single combination was tested to examine the robustness of the method. (**E**) Comparative analysis of antigen-specific frequencies within full combination, single combination and combination minus one. Bar plots represent the mean of 3 independent experiments ± SEM. Full combination conditions are similar to single combinations, and limited background is detected when a given combination is not added.

**Figure S2.**
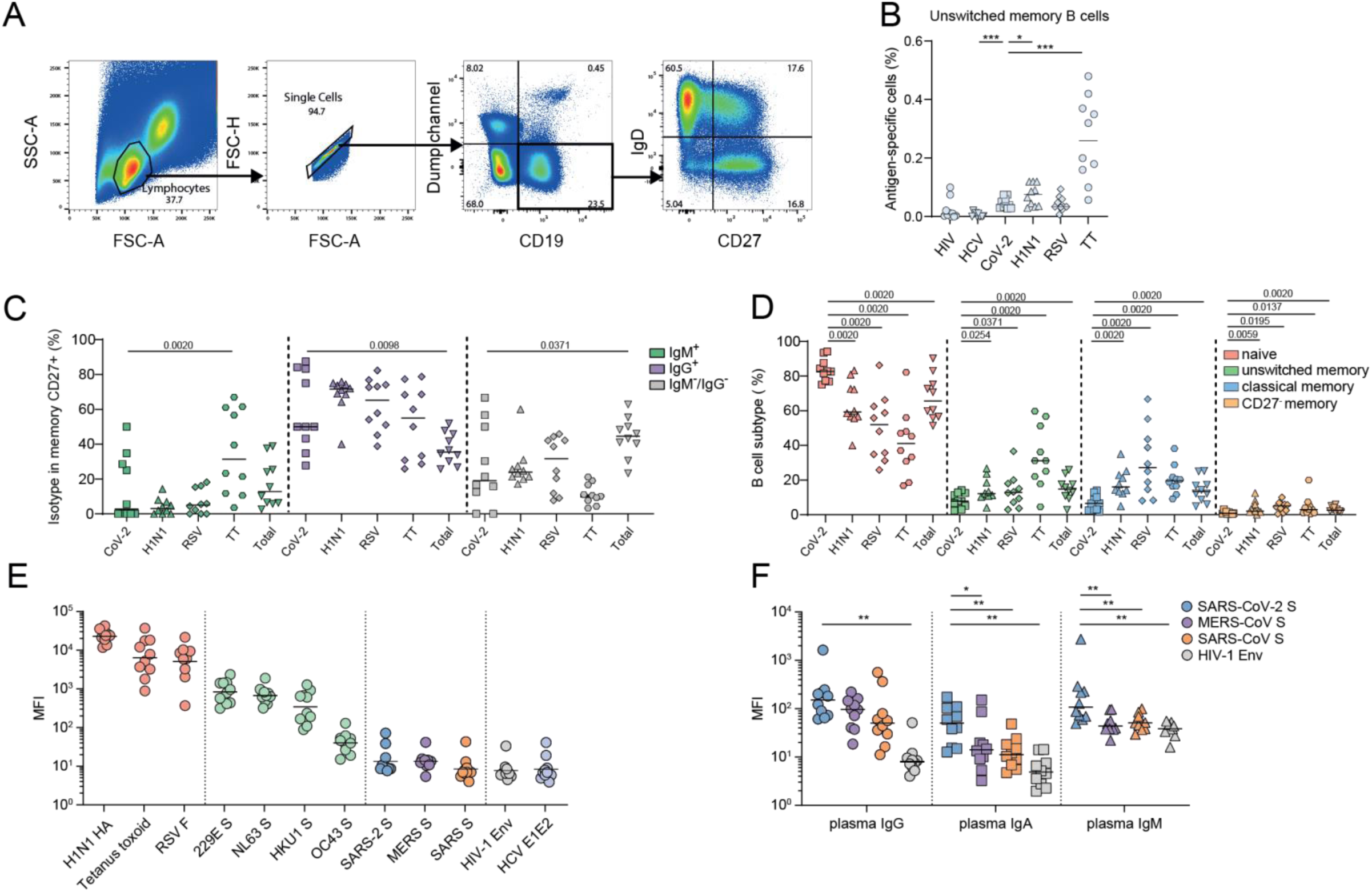
Phenotypic characterization of SARS-CoV-2 S-reactive B cells in unexposed individuals extra data. **(A)** Gating strategies to identify live B cells (CD19^+^ Via^-^ CD14^-^) and define B cell subsets according IgD and CD27 expression in total B cells (IgD^+^/CD27^-^, naive; IgD^+^/CD27^+^, unswitched IgD^+^ memory; IgD^-^/CD27^-^, CD27^-^ memory; IgD^-^/CD27^+^, classical memory). **(B-D)** Statistical differences were tested only in comparison to SARS-CoV-2 condition **(B)** Analysis of frequency of antigen-specific B cells in the Unswitched memory B cells population. **(C)** Analysis of frequency of IgG^+^, IgM^+^, or IgM^-^/IgG^-^ in antigen-specific or Total classical memory B cells. **(D)** Analysis of frequency of antigen-specific B cells in the different B cell subsets (naive, unswitched memory, CD27-memory, classical memory). (**E**) MFI of plasma IgG binding (1:10.000 dilution) for each antigen as measured by custom Luminex assay. (**F**) MFI of plasma IgG, IgA and IgM binding (1:10 dilution) for each antigen as measured by custom Luminex assay.

**Figure S3.**
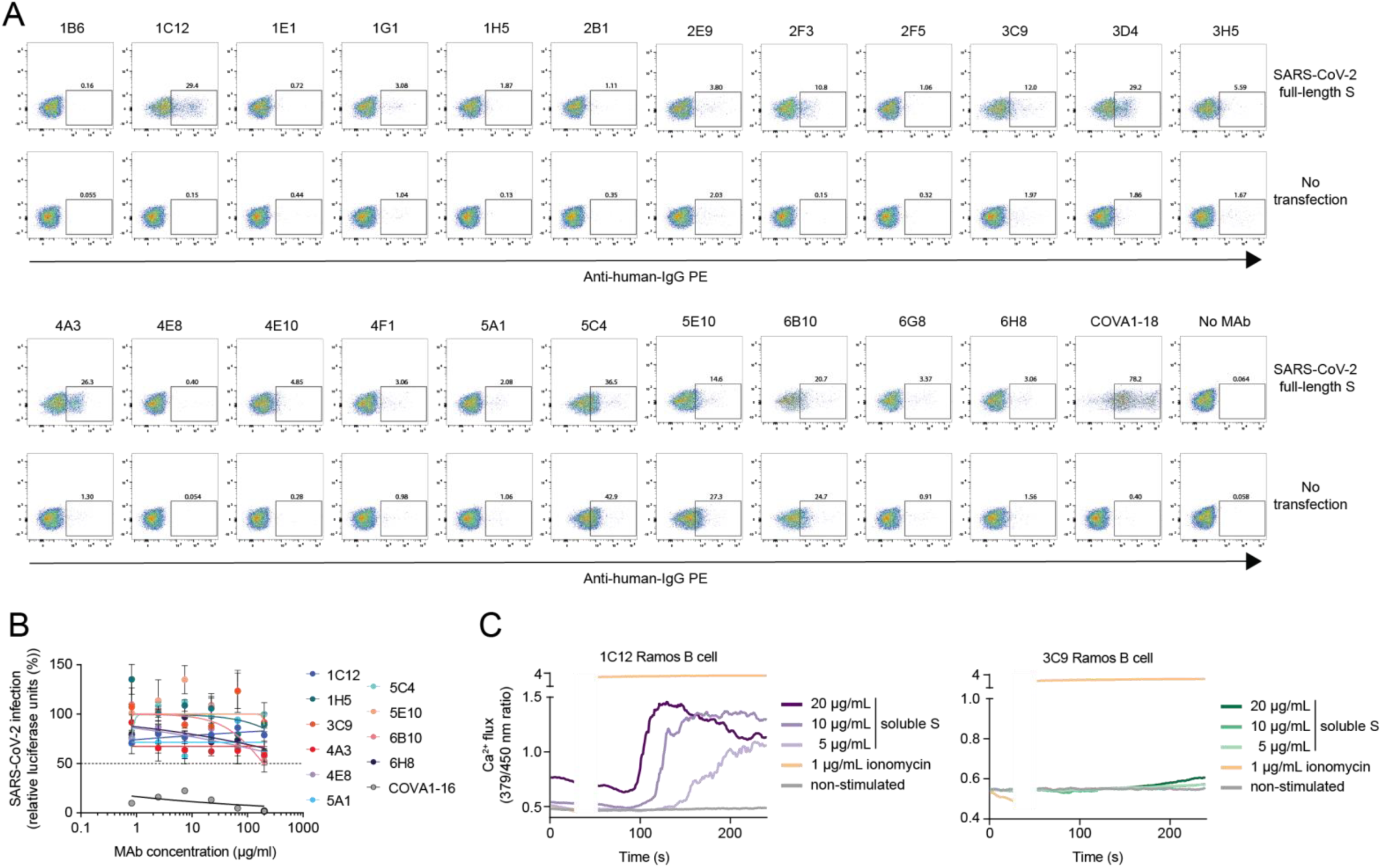
Molecular characterization of MAbs isolated from HDs. (**A**) FACS plots depicting binding to full length SARS-CoV-2 S-transfected or untransfected HEK293T cells for 23 selected MAbs isolated from unexposed individuals and control MAb COVA1-18 isolated from a convalescent COVID-19 patient. **(B)** Neutralization of SARS-CoV-2 pseudovirus of 10 selected MAbs isolated from unexposed individuals and control MAb COVA1-16 isolated from a convalescent COVID-19 patient. (**C**) Ramos B cell activation of 1C12 B cells (top panel) and 3C9 B cells (bottom panel) as measured by calcium (Ca^2+^) flux assay. A baseline without antigen was established between 0 and 30 seconds, after which the measurement was interrupted to add the antigen to the B cells (30-50 seconds). Ionomycin was used at 1 μg/mL as positive control.

**Figure S4.**
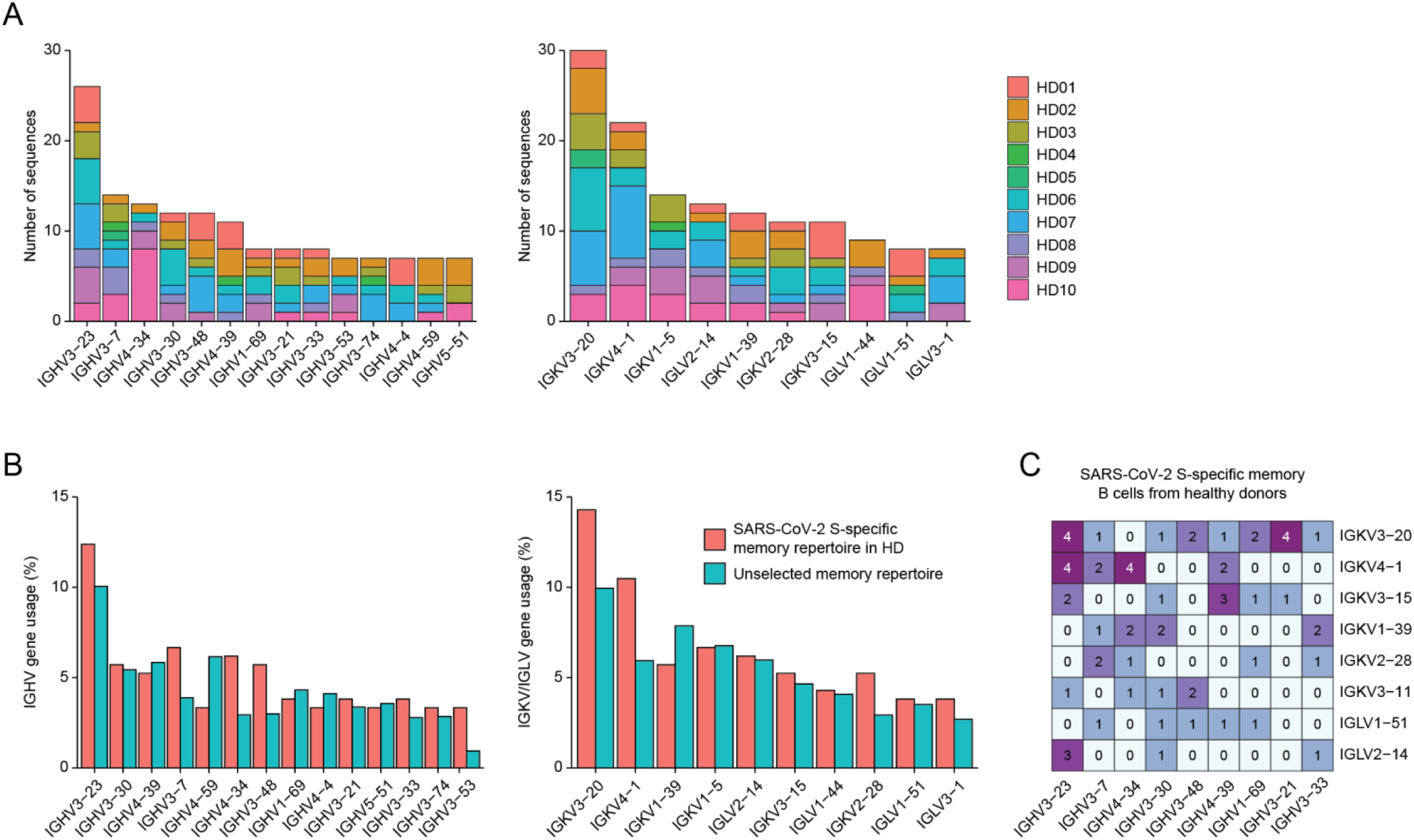
BCR sequence analysis of memory B cells from HDs. (**A**) Number of sequences in the memory cluster as defined in Fig. 3A for each respective IGHV and IGKV/IGLV gene. Each color corresponds to an individual HD. (**B**) IGHV (left panel) and IGKV/IGLV (right panel) gene usage in all recovered sequences in the memory cluster as defined in Fig. 3A (red) and an unselected memory repertoire from (DeKosky et al. 2016). (**C**) Matrix showing the number of pairs with a certain IGHV (x-axis) and IGKV (y-axis) recovered from SARS-CoV-2 S-reactive memory B cells from HDs.

**Figure S5.**
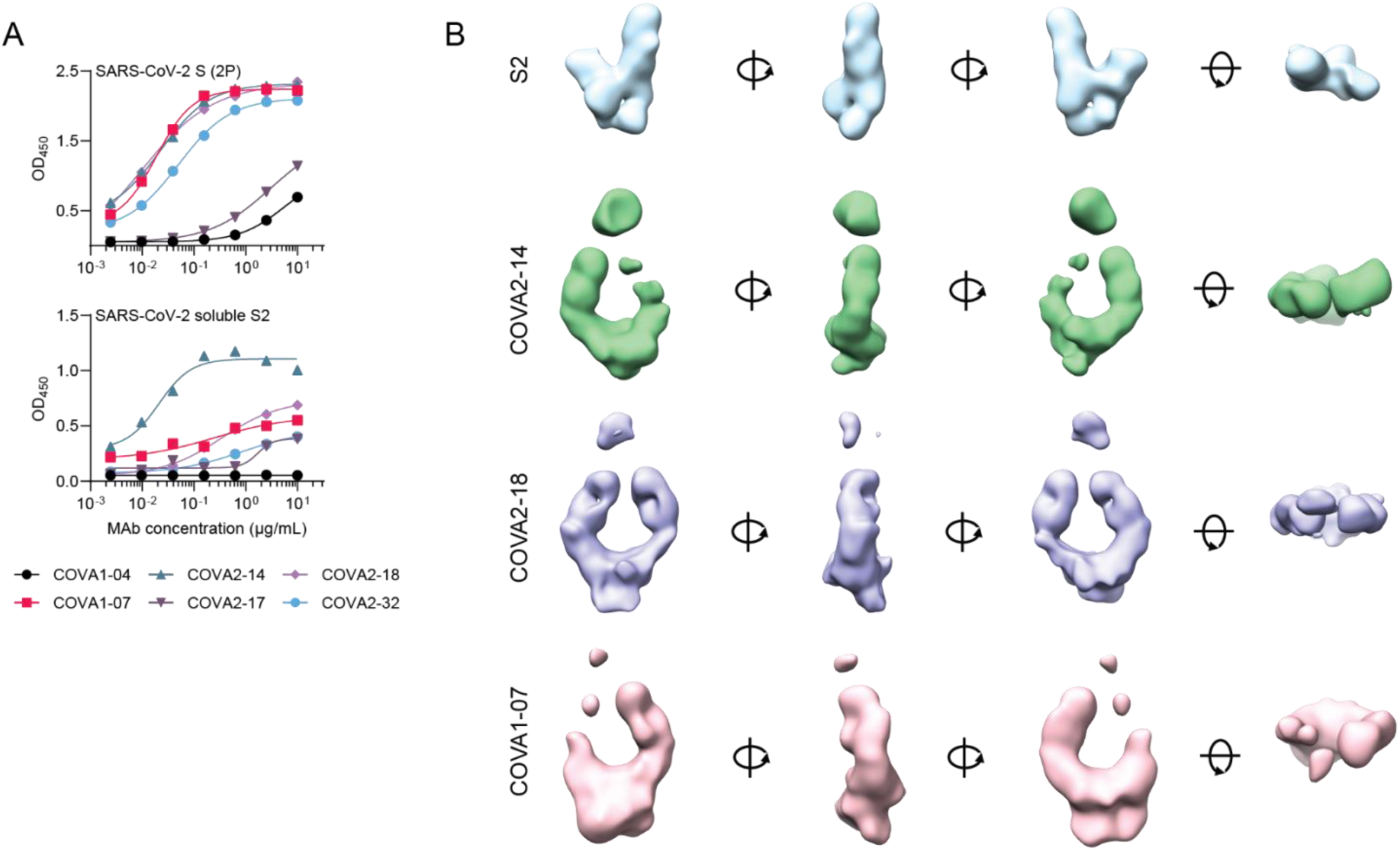
Characterization of IGHV1-69/IGKV3-11 MAbs from convalescent patients. (**A**) Enzyme-linked immunosorbent assay (ELISA) showing binding of six IGHV1-69/IGKV3-11 MAbs isolated from convalescent patients to SARS-CoV-2 S-2P (top panel) and soluble SARS-CoV-2 S2 (bottom panel). (**B**) 3D reconstruction SARS-CoV-2 S2 (top row) and COVA2-14, COVA2-18 and COVA1-07 MAbs in complex with SARS-CoV-2 S2.

**Figure S6.**
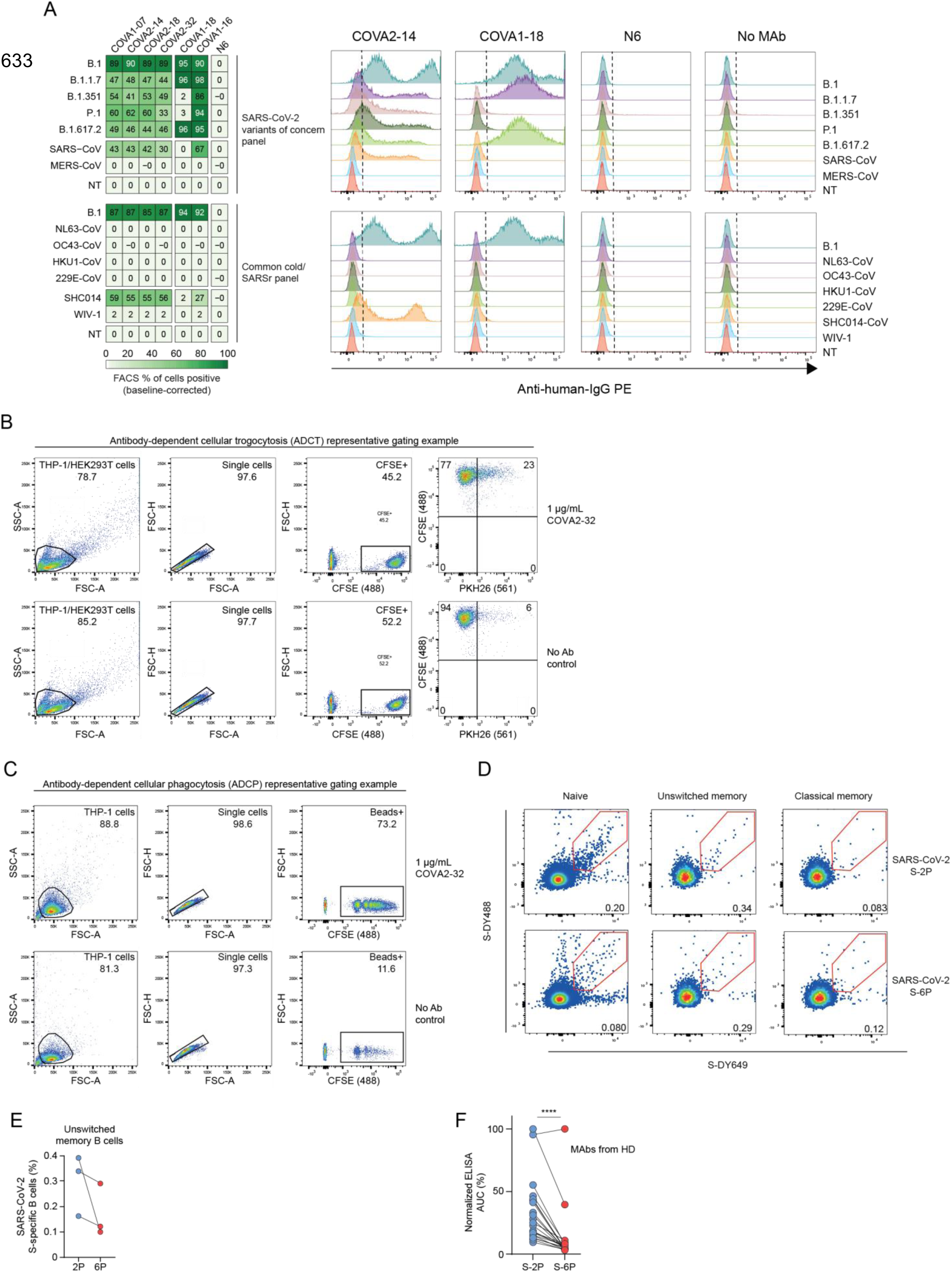
Functional characterization of IGHV1-69/IGKV3-11 MAbs. **(A)** Matrix showing flow cytometric binding assay to SARS-CoV-2 S-transfected or untransfected HEK293T cells (NT) for IGHV1-69/IGKV3-11 MAbs (COVA1-07, COVA2-14, COVA2-18 and COVA2-32), anti-SARS-CoV-2 MAbs isolated from convalescent patients (COVA1-18 and COVA1-16) and control anti-HIV-1 MAb N6. Numbers and colors in the boxes represent the percentage of cells showing binding to a particular MAb. Representative MFI peaks are shown for COVA2-14, COVA1-18, N6 and no MAb are shown on the right. (**B-C**) Representative gating strategy for antibody-dependent cellular trogocytosis (ADCT, (**B**)) and antibody-dependent cellular phagocytosis (ADCP, (**C**)). The top panels show an example of COVA2-32, the bottom panels show the no MAb controls. (**D**) Representative gating strategy showing binding of naive (left), unswitched memory (middle) and classical memory (right) B cells to SARS-CoV-2 S-2P in two colors (top row) and SARS-CoV-2 S-6P in two colors (bottom row). (**E**) Connected dot plots showing the frequency of unswitched B cells of SARS-CoV-2 S-2P- or S-6P-specific B cells (%) in three HDs. (**F**) Connected dot plot showing the ELISA area under the curve (AUC) of isolated MAbs from HDs to SARS-CoV-2 S-2P- or S-6P. ****, *p* < 0.0001.

